# ISWI chromatin remodeling complexes recruit NSD2 and H3K36me2 in pericentromeric heterochromatin

**DOI:** 10.1101/2023.10.20.563387

**Authors:** Naoki Goto, Kazuma Suke, Nao Yonezawa, Hidenori Nishihara, Yuko Sato, Tomoya Kujirai, Hitoshi Kurumizaka, Kazuo Yamagata, Hiroshi Kimura

**Affiliations:** School of Life Science and Technology, Tokyo Institute of Technology; Faculty of Biology-Oriented Science and Technology, Kindai University; Department of Advanced Bioscience, Graduate School of Agriculture, Kindai University; Cell Biology Center, Institute of Innovative Research, Tokyo Institute of Technology; Instite of Quantitative Biology, University of Tokyo

## Abstract

Histone H3 lysine36 dimethylation (H3K36me2) is generally distributed in the gene body and euchromatic intergenic regions. However, we found that H3K36me2 is enriched in pericentromeric heterochromatin in some mouse cell lines. We here revealed the mechanism of heterochromatin targeting of H3K36me2. Among several H3K36 methyltransferases, NSD2 was responsible for inducing heterochromatic H3K36me2. Depletion and overexpression analyses of NSD2-associating proteins revealed that NSD2 recruitment to heterochromatin was mediated through the imitation switch (ISWI) chromatin remodeling complexes, such as BAZ1B-SMARCA5 (WICH), which directly binds to AT-rich DNA via a BAZ1B domain containing AT-hook-like motifs. The abundance and stoichiometry of NSD2, SMARCA5, and BAZ1B or BAZ2A could determine the localization of H3K36me2 in different cell types. To explore the physiological role of heterochromatic H3K36me2, we analyzed mouse tissues and embryos. As a result, H3K36me2 was found in heterochromatin at the 2- to 4-cell stages of mouse preimplantation embryos, suggesting its involvement in developmental regulation.

**Summary:** The authors discovered histone H3K36me2, which is believed to be enriched in potentially active genomic regions, is also located in transcriptionally inactive regions called heterochromatin in some cell types. The detailed molecular mechanism of its heterochromatin targeting is now revealed.

## Introduction

In eukaryotic cells, DNA is packed into chromatin, which could be categorized into highly condensed heterochromatin and loosely decondensed euchromatin (Woodcock and Ghosh, 2010). Pericentromeric heterochromatin (PCH) is composed of repeat elements, including α-satellites in humans (Altemose et al., 2022) and major satellites in mice (Saksouk et al., 2015). Except during specific stages of development and under some cellular stresses, transcription from PCH is mostly repressed (Smurova and Wulf, 2018). In mouse interphase cell nuclei, PCH forms microscopically visible clusters termed chromocenters. PCH in mice is epigenetically characterized by a high level of histone H3 lysine 9 trimethylation (H3K9me3), H4 lysine 20 trimethylation (H4K20me3), cytosine methylation on CpG sites, and low histone acetylation (Probst and Almouzni, 2008). Other histone modifications, such as H3 lysine 27 monomethylation (H3K27me1) (Peters et al., 2003), H3 lysine 56 trimethylation (H3K56me3) (Jack et al., 2013), H3 lysine 64 trimethylation (H3K64me3) (Lange et al., 2013), and H3 lysine 79 trimethylation (H3K79me3) (Ooga et al., 2008), have been reported to localize to PCH, but the detailed molecular mechanism for specialized localization and physiological significance, are not well understood.

In mammals, histone H3 lysine36 di- and tri-methylation (H3K36me2 and H3K36me3) are known to function in euchromatin, methylated by histone methyltransferases such as nuclear receptor SET domain-containing 1/2/3 (NSD1/2/3) and SETD2 (Wagner and Carpenter, 2012) and demethylated by lysine demethylases such as KDM2A/JHDM1A (Blackledge et al., 2010). H3K36me2 serves as a marker to attract DNMT3A to the intergenic region and contributes to the establishment of DNA methylation patterns (Weinberg et al., 2019).

H3K36me3 accumulates in transcriptionally active gene bodies by Set2 (an ortholog of SETD2) in yeast and by SETD2 in mouse ES cells, serving as a scaffold for the histone deacetylation complex RPD3 (Carrozza et al., 2005; Guan et al., 2023) and DNA methyltransferase DNMT3B (Neri et al., 2017) to repress aberrant transcription. Such regulation of DNA methylation via H3K36me2/3 has been shown to play an important role in germline development (Xu et al., 2019; Shirane et al., 2020). H3K36me2 also shows antagonistic distribution with H3K27me3. In mouse ES cells, H3K36me2 protects the genome from excessive accumulation of PRC2-dependent H3K27me3 (Streubel et al., 2018; Chen et al., 2022). Similar antagonism has been observed in multiple myeloma, which highly express NSD2 (Popovic et al., 2014).

In addition to the function of H3K36me2/3 in euchromatin, several studies have reported their roles in heterochromatin. H3K36me2 is enriched in the pericentromere regions of Drosophila eye discs and represses transcription of transposons such as long interspersed nuclear element (LINE) and long-terminal repeat (LTR) (Chaouch et al., 2021). KDM2A depletion causes H3K36me2 enhancement and increased transcription of satellite repeats in the PCH of mouse and human cells (Frescas et al., 2008). H3K36me3 accumulation in PCH has also been observed in mouse ES cells (Chantalat et al., 2011). In addition, H3K9me3 and H3K36me3 co-localize in some intergenic regions and gene bodies in mouse ES cells and can function as potential enhancers (Barral et al., 2022). In fission yeast, H3K36me3 by Set2 is required for histone deacetylation for RNAi-dependent heterochromatin establishment (Chen et al., 2008), maintenance of condensed chromatin near subtelomeres (Matsuda et al., 2015), and transcriptional repression of pericentromeres and subtelomeres (Georgescu et al., 2020).

ATP-dependent chromatin remodeling complexes are widely conserved in eukaryotes. Higher eukaryotes have four subfamilies: switch/sucrose non-fermenting (SWI/SNF), chromodomain-helicase DNA-binding (CHD), imitation switch (ISWI), and inositol-requiring 80 (INO80) (Hasan and Ahuja, 2019). These complexes utilize ATP hydrolysis-derived energy to drive nucleosome assembly, nucleosome sliding, and histone exchange to regulate transcription, replication, and repair (Clapier et al., 2017). The CHD and ISWI complexes play a particularly crucial role in heterochromatin. The NuRD complex recognizes H3K9me3 via the PHD domain of CHD4 (Musselman et al., 2012) and methylated DNA via the MBD2 subunit (Zhang et al., 1999), suggesting that the complex is involved in heterochromatin regulation and transcriptional repression. ISWI subfamily containing SMARCA5/SNF2H as an ATPase consists of five members in higher eukaryotes; CHRAC (with BAZ1A/ACF1, CHRAC15, and CHRAC17), ACF (with BAZ1A), WICH (with BAZ1B/WSTF), NoRC (with BAZ2A/TIP5), and RSF (with and RSF1) (Bartholomew, 2014). ACF accumulates in PCH during the S phase of mouse cells, and depletion of BAZ1A delays late S phase progression, suggesting that ACF seems to contribute to PCH replication (Collins et al., 2002). WICH is associated with DNA replication foci during the S phase in mouse cells in a PCNA-dependent manner and concentrated on PCH in the late S phase. Depletion of each subunit of WICH increases H3K9me3 and HP1α/β levels and reduces chromatin accessibility after replication (Poot et al., 2004). It has also been reported that WICH localizes to inactive X chromosomes in the late S phase in human cells (Culver-Cochran and Chadwick, 2012). NoRC suppresses rRNA transcription by inducing DNA methylation and histone deacetylation on the rDNA promoter in the nucleolus (Zhou et al., 2002; Santoro et al., 2002). In addition, BAZ2A depletion in mouse cells causes genomic instability by decreasing H4K20me3 and H3K9me3 in major satellite repeats and rDNA (Guetg et al., 2010). ISWI complexes are thought to be differentially recruited to their target regions by SMARCA5 partner factors that bind to different histone modifications (Ito et al., 1999; Ribeyre et al., 2016; Strohner et al., 2001; Li et al., 2006; Tallant et al., 2015); however, the mechanism of target recognition on PCH is not well understood.

In a series of immunofluorescence studies using mouse cells, we noticed that some cells exhibit H3K36me2 enrichment in PCH while others show typical euchromatic distributions. This intriguing observation led us to investigate the mechanism regulating H3K36me2 localization. Here we show that H3K36me2 could be localized to PCH via the ISWI-NSD2 axis.

## Results

### H3K36me2 is enriched in pericentromeric heterochromatin in several mouse cell lines

To investigate the nuclear distribution of H3K36me2 in various mouse cell types, we immunostained H3K36me2 with a specific mouse monoclonal antibody (clone 2C3/CMA332; Rechtsteiner et al., 2010) and detected it using a confocal microscope. H3K9me3 was co-stained as a marker of constitutive heterochromatin, or chromocenter. In Pmi28 and C2C12 myoblasts and MC12 embryonic carcinoma cells, H3K36me2 showed typical euchromatin distribution (Fig. 1A and S1A). In contrast, in immortalized mouse embryonic fibroblasts (iMEFs), NIH3T3 fibroblasts, and embryonic stem cells (ESCs), H3K36me2 was concentrated on PCH where H3K9me3 is enriched (Fig. 1B and S1A). Intensity measurements confirmed that the enrichment of H3K36me2 in PCH was higher in iMEFs, NIH3T3, and ESCs, compared to the other three cell lines (Fig. 1C). Using immunofluorescence with another H3K36me2-specific antibody (rabbit monoclonal clone EPR16994(2); Bressan et al., 2021), we observed the same localization patterns (Fig. S1B). This indicates that the variations in H3K36me2 distribution are not due to antibody staining artifacts. In iMEFs that exhibit H3K36me2 PCH localization, other histone marks such as H3K27me3, H3K9me2, and H3K36me3, were not concentrated in PCH (Fig. S1C-E), suggesting a unique feature of H3K36me2 independently of these histone marks.

**Figure 1.**
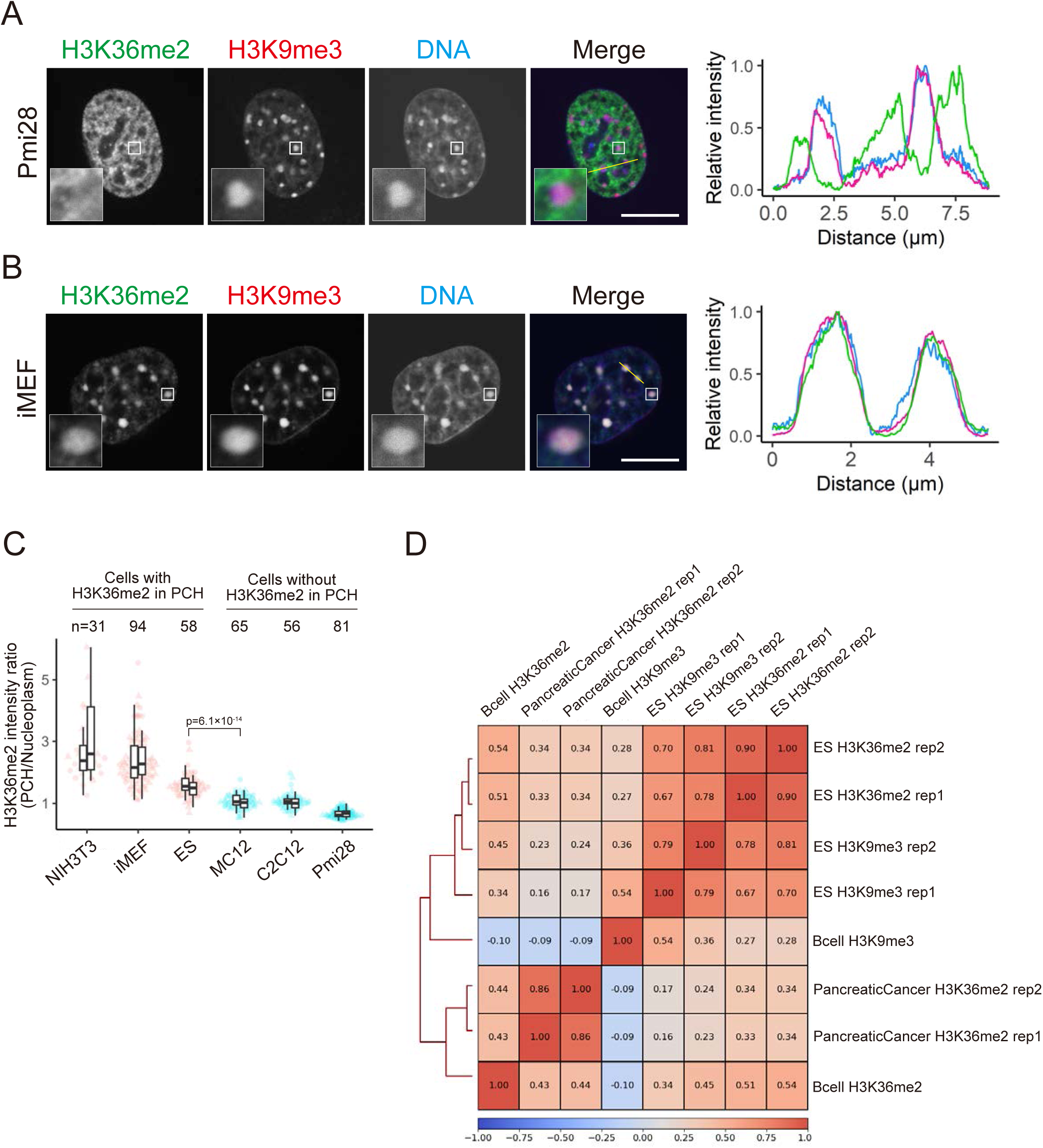
H3K36me2 is enriched in pericentromeric heterochromatin in some mouse cells. (A-C) Localization of H3K36me2 by Immunofluorescence. Pmi28 cells (A) and iMEFs (B) were fixed and stained with anti-H3K36me2 (green), anti-H3K9me3 (red), and Hoechst 33342 (blue). (A and B) Single optical sections of confocal microscope images are shown. Insets show magnified views of indicated areas. Line plots indicate intensity profiles of H3K36me2, H3K9me3, and DNA along the yellow line in the merged images. (C) Quantification of the intensity ratio of H3K36me2 in PCH to that in the nucleoplasm in various mouse cell lines. Two box plots display data from two biological replicates with dots representing individual cells (round for the first experiment and triangular for the second). Center lines represent medians; box limits indicate the 25th and 75th percentiles; whiskers extend 1.5 times the interquartile range from these percentiles. The numbers of total cells analyzed (n) and p-value comparing ESCs (H3K36me2 in PCH) to MC12 cells (H3K36me2 in euchromatin), calculated using the Mann-Whitney U test, are indicated. (D) Genome-wide correlation heatmap of H3K36me2 and H3K9me3 in mouse ESCs, B cells, and pancreatic cancer cells. Spearman’s correlation coefficients are indicated inside cells. Scale bars, 10 μm.

To validate the immunofluorescence data, we analyzed the genome-wide distribution of H3K36me2 in different mouse cell types using the public ChIP-seq datasets (Barral et al., 2022; Vian et al., 2018; Yuan et al., 2020). The correlation analysis showed that H3K36me2 and H3K9me3 were highly correlated in ESCs, but not in B cells and pancreatic cancer cells (Fig. 1D). H3K36me2 in B cells and pancreatic cancer cells exhibited anti-correlation with H3K9me3 in B cells. Upon analyzing enrichments in repeat elements with transcription units, H3K9me3 was enriched in LINE and LTR, particularly at the transcription start site (TSS), in both ESCs and B cells (Fig. S1F and S1G). This is consistent with the silencing role of this modification in those repeats. H3K36me2 in ESCs, but not in B cells, was also enriched in LINE and LTR, reflecting its genome-wide correlation with H3K9me3. H3K36me2 in both ESCs and B cells was enriched at TSS of SINE, which is rich in CpG that can be methylated (Varshney et al., 2015), in line with the role of H3K36me2 in establishing DNA methylation in euchromatin (Weinberg et al., 2019). Thus, the enrichment of H3K36me2 in H3K9me3-rich genomic regions in ESCs observed in public datasets well agreed with immunofluorescence results.

### H3K9me3 and DNA methylation are not required for H3K36me2 in pericentromeric heterochromatin

H3K9me3 and DNA methylation are the major epigenetic marks concentrated in PCH in mouse cells. To examine if these marks are required for PCH H3K36me2 localization, we used knockout (KO) cell lines lacking H3K9me3 and DNA methylation. 5KO iMEFs lack H3K9me3 because five H3K9me2/3-methyltransferases, i.e., Suv39h1, Suv39h2, Setdb1, G9a, and Glp, are deleted (Fukuda et al., 2023). TKO ESCs lack DNA methylation because three DNA methyltransferases, i.e., Dnmt1, Dnmt3a, and Dnmt3b, are deleted (Tsumura et al., 2006). In both of these cell lines, H3K36me2 remained enriched in PCH (Fig. S2A and S2B), although the enrichment was slightly reduced in 5KO iMEFs. Therefore, the localization of H3K36me2 in PCH does not require H3K9me3 and DNA methylation, and the related methyltransferases.

### NSD2 is responsible for H3K36me2 in pericentromeric heterochromatin

To identify which H3K36 methyltransferase targets PCH and mediates H3K36me2, we analyzed the localization of transiently expressed Halo-tagged enzymes in NIH3T3 cells (Fig. S2C). Among several methylation and demethylation enzymes, NSD2/WHSC1 and KDM2A/JHDM1A localized to PCH. KDM2A is indeed known to localize to PCH in an HP1-dependent manner (Borgel et al., 2017). Although H3K9me3 is required for HP1 localization to the PCH (Lachner et al., 2001), H3K36me2 was still concentrated in H3K9me3-deficient cells (Fig. S2A), suggesting that KDM2A per se does not play a primary role in H3K36me2 enrichment in PCH. Therefore, we focused on NSD2-mediated H3K36me2 regulation. Hereafter, we used iMEFs and Pmi28 cells for cells exhibiting H3K36me2 in PCH and euchromatin, respectively, for overexpression and iMEFs for KO studies.

Overexpressed Halo-NSD2 localized to PCH in iMEFs but not obvious in Pmi28 cells (Fig. 2A and 2B). Measuring PCH-to-nucleoplasm intensity ratios in individual nuclei revealed that Halo-NSD2 was enriched in PCH in all iMEF nuclei but its enrichment was lower in Pmi28 nuclei (Fig. 2C). In iMEFs, Halo-NSD2 expression did not affect the enrichment of H3K36me2 in PCH (Fig. 2D), suggesting that increased levels of NSD2 had little effect. In Pmi28 cells, when Halo-NSD2 was expressed, PCH H3K36me2 was induced albeit at lower levels compared to iMEFs, suggesting that NSD2 alone is not sufficient for inducing PCH H3K36me2.

**Figure 2.**
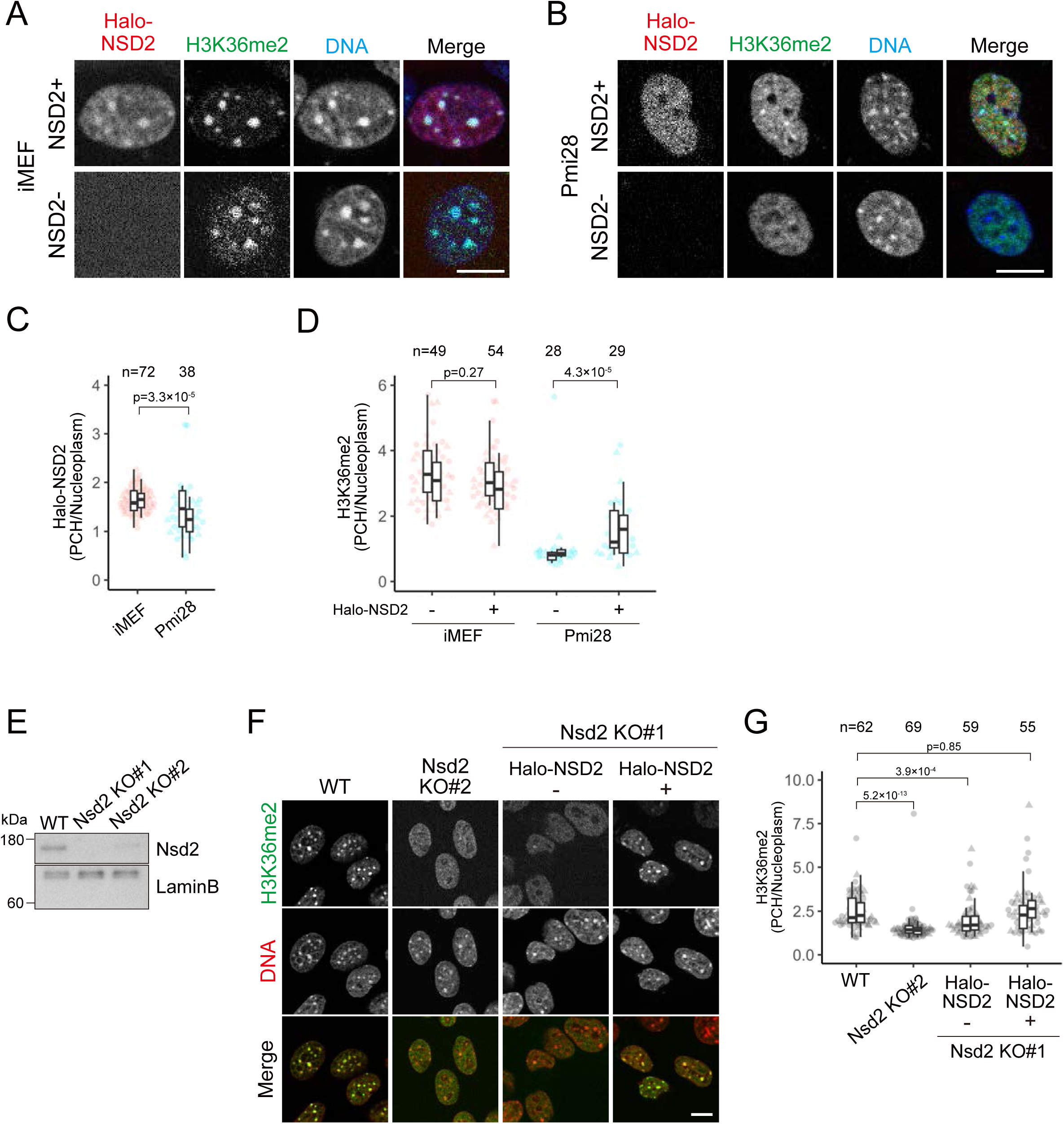
NSD2 is the enzyme responsible for H3K36me2 at pericentromeric heterochromatin. (A-D) Effect of Halo-NSD2 expression on H3K36me2 localization. iMEFs and Pmi28 cells were transfected with the Halo-NSD2 expression vector, and treated with JF646 HaloTag ligand and Hoechst33342. After live imaging, cells were fixed and stained with anti-H3K36me2. (A and B) Examples of single optical sections for Halo-NSD2 (red), H3K36me2 (green), and Hoechst33342 (blue). Halo-NSD2 expressed (NSD2+) and unexpressed (NSD2-) iMEFs (A) and Pmi28 cells (B) are shown. (C and D) PCH-to-nucleus intensity ratios for Halo-NSD2 in live cells (C) and H3K36me2 by immunofluorescence (D). (E-G) Effect of NSD2 KO on H3K36me2 localization. (E) The protein levels of NSD2 evaluated by Western blotting in WT and NSD2 KO#1 and KO#2 iMEFs. Lamin B served as a loading control. Positions of size standards are indicated on the left. (F and G) Immunofluorescence and quantification. (F) WT, NSD2 KO#2, and KO#1 without and with Halo-NSD2 were stained with anti-H3K36me2 (green) and Hoechst33342 (red). Single optical section images are displayed. (G) PCH-to-nucleus intensity ratios for H3K36me2. See Fig. 1 legend for the details of box plots. Scale bars, 10 μm.

To validate the role of NSD2 in PCH H3K36me2 in iMEFs, we generated NSD2 KO cell lines using the CRISPR/Cas9 system (Fig. 2E). NSD2 depletion reduced H3K36me2 levels in PCH, and this phenotype was rescued by Halo-NSD2 expression (Fig. 2F and 2G). Thus, this result indicates that NSD2 is responsible for inducing H3K36me2 in PCH.

### SMARCA5 is required for NSD2 localization and H3K36me2 in PCH

As the PWWP domain of NSD2 binds to H3K36me2 and stabilizes NSD2 on chromatin (Sankaran et al., 2016), it is possible that once H3K36me2 is introduced into PCH by ectopic NSD2 expression, the modification could be maintained through positive feedback. However, the mechanism of how NSD2 initially targets remains uncertain. Therefore, we searched for a potential NSD2 interactor on heterochromatin. Among the NSD2-interacting factors comprehensively analyzed in a previous report (Huang et al., 2019), there were several candidate proteins associated with PCH, including SMARCA5, PARP1, ORC2, MKI67, MECP2, KDM2A, and HMGB1 (Fig. 3A) (Malla et al., 2023; Quenet et al., 2008; Prasanth et al., 2010; Sobecki et al., 2016; Nan et al., 2007; Frescas et al., 2008; Pallier et al., 2003). These candidate factors and NSD2 were knocked down in iMEFs using the lentivirus shRNA expression system, and the number of cells clearly showing PCH H3K36me2 was counted (Fig. 3B). The knockdown efficiency was verified by qPCR. Knockdown of NSD2 had the most drastic effect, reassuring the critical role of NSD2 in PCH H3K36me2. Among the candidate proteins, knockdown of the chromatin remodeling factor SMARCA5/SNF2H substantially reduced the number of cells exhibiting PCH H3K36me2. Therefore, we performed a deeper analysis of SMARCA5.

**Figure 3.**
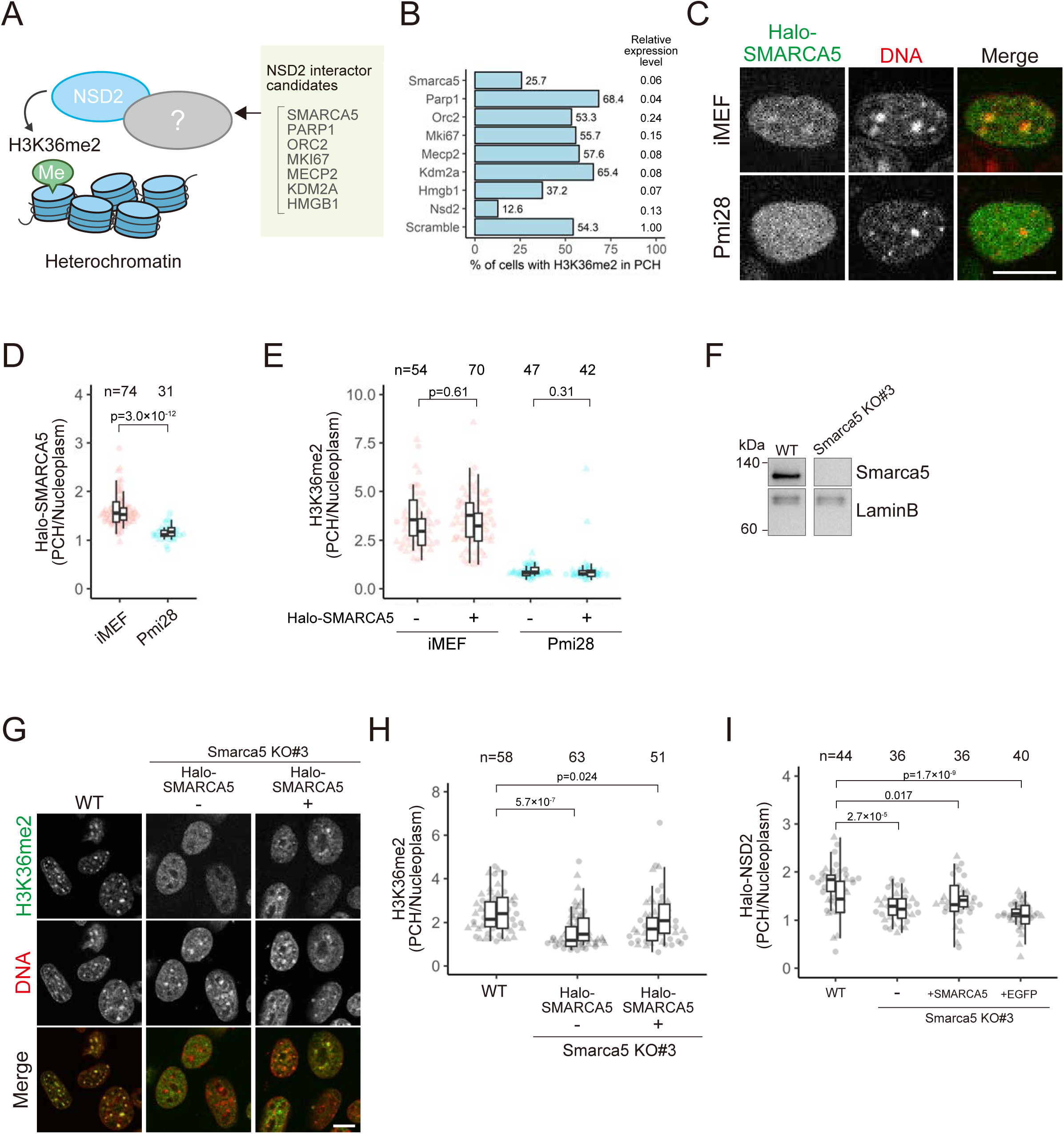
SMARCA5 modulates NSD2 localization and H3K36me2 levels in PCH. (A) Candidates of NSD2-interacting proteins that mediate heterochromatic H3K36me2. Among the NSD2-interacting factors previously identified (Huang et al., 2019), some are known to be enriched in PCH. (B) Percentages of cells clearly showing PCH H3K36me2 when the indicated proteins are depleted by shRNA-mediated knockdown. The knockdown efficiencies are indicated on the right as relative expression levels to the control (scramble) shRNA, measured by RT-qPCR. (C-E) Localization of Halo-SMARCA5 and H3K36me2. iMEFs and Pmi28 cells, transfected with the Halo-SMARCA5 expression vector, were incubated in JF646 HaloTag ligand and Hoechst33342. After live imaging, cells were fixed and stained with anti-H3K36me2. (C) Single optical sections for Halo-SMARCA5 (green) and Hoechst33342 (red) in living cells. (D and E) PCH-to-nucleus intensity ratios of Halo-SMARCA5 in live cells (D) and H3K36me2 by immunofluorescence (E). (F-I) Effects of SMARCA5 KO on the localization of H3K36me2 and NSD2. (F) SMARCA5 protein levels evaluated by Western blotting in WT and SMARCA5 KO#3 iMEFs. Lamin B served as a loading control. The positions of size standards are indicated on the left. (G and H) Immunofluorescence of H3K36me2 and quantification. (G) WT and SMARCA5 KO#3 without and with Halo-SMARCA5 were stained with anti-H3K36me2 (green) and Hoechst33342 (red). Single optical section images are shown. (H) PCH-to-nucleus intensity ratios of H3K36me2. (I) Intensity ratios of Halo-NSD2 in WT and SMARCA5 KO#3 cells without and with SMARCA5-sfGFP or EGFP. See Fig. 1 legend for the details of box plots. Scale bars, 10 μm.

Firstly, we overexpressed Halo-SMARCA5 in iMEFs and Pmi28 cells to examine the localization of the expressed protein and H3K36me2. Halo-SMARCA5 was concentrated in PCH in iMEFs but not much in Pmi28 cells (Fig. 3C and 3D). SMARCA5 expression did not affect PCH H3K36me2 levels in both iMEFs and Pmi28 cells (Fig. 3E). These data suggest that SMARCA5 alone is not sufficient to target PCH and induce H3K36me2 there.

Secondly, we established SMARCA5 KO iMEFs to investigate the requirement of SMARCA5 on H3K36me2 and NSD2 localization in PCH (Fig. 3F). The loss of SMARCA5 led to a decrease in PCH H3K36me2 levels, and this phenotype was rescued to some extent by the expression of Halo-SMARCA5 (Fig. 3G and 3H). Furthermore, in SMARCA5 KO cells, PCH localization of transfected Halo-NSD2 was lower, and the expression of SMARCA5-sfGFP, but not EGFP alone, also partially restored PCH H3K36me2 enrichment (Fig. 3I). Taken together, we concluded that SMARCA5 is required, but not sufficient, for PCH H3K36me2 by recruiting NSD2.

### BAZ1B regulates PCH localization of SMARCA5, NSD2, and H3K36me2

In mammalian cells, SMARCA5 forms ISWI complexes with various partner factors, such as ACF with BAZ1A, WICH with BAZ1B, and NoRC with BAZ2A (Bartholomew, 2014). Chromatin remodeling is driven by SMARCA5 through its ATPase activity, while the partner factors are thought to modulate the activity and recruitment of the ISWI complexes to target sites (Ito et al., 1999; Ribeyre et al., 2016; Strohner et al., 2001). Both BAZ1A and BAZ1B have been reported to accumulate at the PCH in the S phase and to promote proper heterochromatin replication (Collins et al., 2002; Poot et al., 2004; Bozhhenok et al., 2002). BAZ2A also associates with pericentromeric repeats and rDNA in the nucleolus and contributes to its genome stabilization (Guetg et al., 2010). Therefore, we hypothesized that BAZ1A, BAZ1B, and/or BAZ2A in the ISWI complexes might bind to PCH to recruit NSD2 (Fig. 4A). To test whether these BAZ proteins target PCH, we expressed their Emerald-green fluorescent protein (emGFP)-tagged versions in iMEFs and Pmi28 cells. Whereas BAZ1A diffused throughout the nucleoplasm in most cells of both cell lines (Fig. S3A), BAZ1B and BAZ2A were concentrated in PCH in both cell types (Fig. 4B, 4C, S3B, and S3C). Overexpression of BAZ1B and BAZ2A however did not induce PCH H3K36me2 in Pmi28 cells (Fig. 4D and S3D), probably because other proteins that are critical for PCH H3K36me2, such as SMARCA5 and NSD2, are not abundantly present.

**Figure 4.**
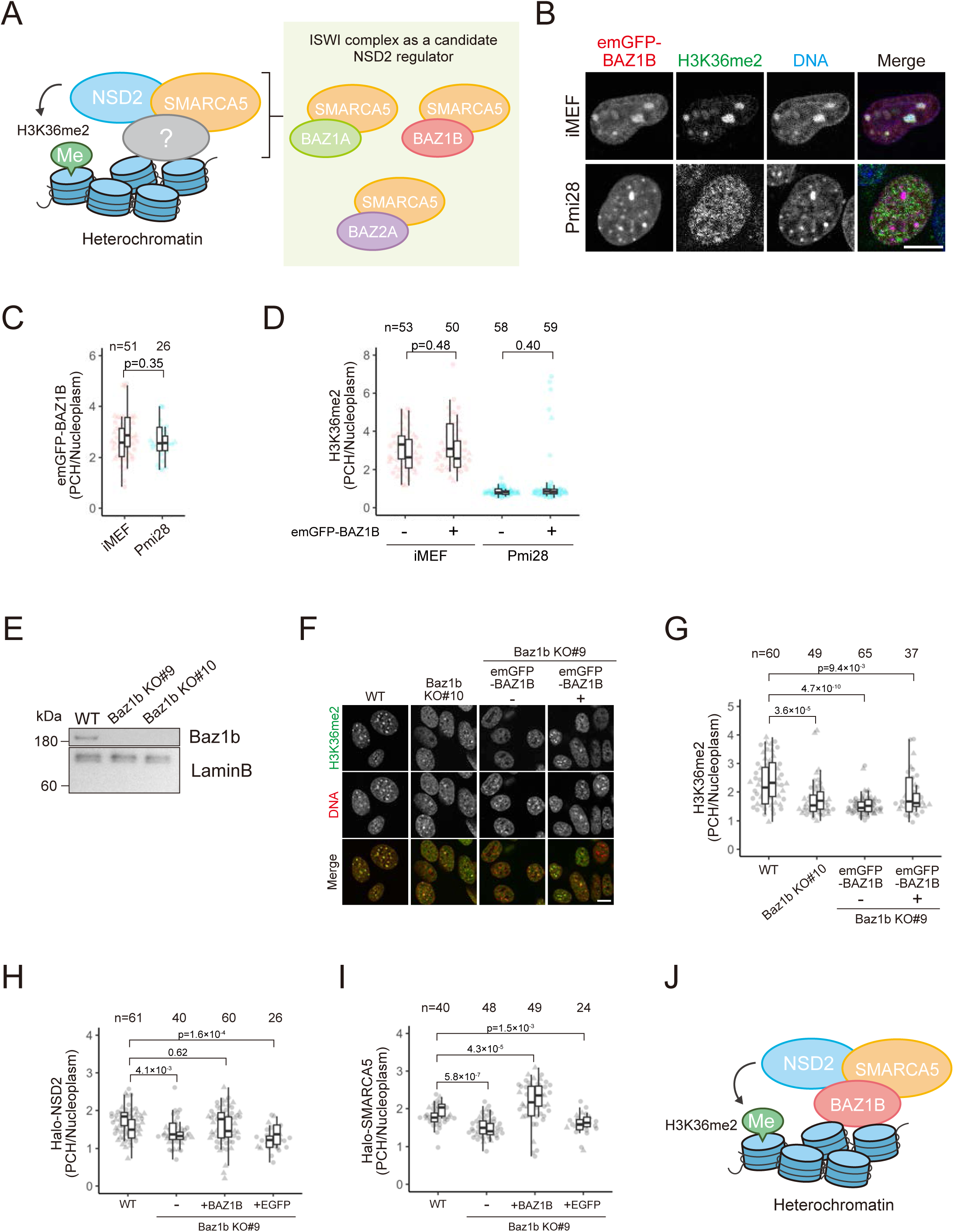
BAZ1B regulates the PCH localization of SMARCA5 and NSD2. (A) Candidates of ISWI complexes that bind to chromatin and recruit NSD2 to PCH. (B-D) Localization of emGFP-BAZ1B and H3K36me2. iMEFs and Pmi28 cells transfected with emGFP-BAZ1B expression vector were stained with Hoechst33342 for live cell imaging. For immunofluorescence, cells were fixed and stained with anti-H3K36me2. (B) Examples of single optical sections for emGFP-BAZ1B (red), H3K36me2 (green), and Hoechst33342 (blue). (C and D) PCH-to-nucleus intensity ratios of emGFP-BAZ1B in living cells (C) and H3K36me2 by immunofluorescence (D). (E-I) Effects of BAZ1B KO on the localization of H3K36me2, NSD2, and SMARCA5. (E) BAZ1B protein levels evaluated by Western blotting in WT and BAZ1B KO#9 and KO#10 iMEFs. Lamin B served as a loading control. The positions of size standards are indicated on the left. (F and G) Immunofluorescence of H3K36me2 and quantification. (F) WT, BAZ1B KO#10, and KO#9 without and with emGFP-BAZ1B were stained with anti-H3K36me2 (green) and Hoechst33342 (red). Single optical section images are shown. (G) PCH-to-nucleus intensity ratios of H3K36me2. (H and I) Intensity ratios of Halo-NSD2 (H) and Halo-SMARCA5 (I) in WT and BAZ1B KO#9 cells without and with emGFP-BAZ1B or EGFP. See Fig. 1 legend for the details of box plots. (J) Schematic drawing of NSD2 recruitment to PCH through the BAZ1B-SMARCA5 complex (WICH). Scale bars, 10 μm.

To understand the role of BAZ1B and BAZ2A in PCH H3K36me2, we attempted to establish single and double KO iMEFs; however, only BAZ1B KO lines were obtained (Fig. 4E) leading us to focus on the function of BAZ1B. PCH H3K36me2 was indeed diminished in BAZ1B KO cells and this phenotype was partially restored by the introduction of emGFP-BAZ1B (Fig. 4F and 4G). The levels of co-expressed Halo-NSD2 and Halo-SMARCA5 at PCH were both decreased in BAZ1B KO cells (Fig. 4H and 4I). The expression of emGFP-BAZ1B, but not EGFP, in KO cells rescued and increased the levels of NSD2 (Fig. 4H) and SMARCA5 (Fig. 4I) in PCH. These results indicate that BAZ1B facilitates PCH localization of NSD2, SMARCA5, and H3K36me2 (Fig. 4J).

### BAZ1B binds to AT-rich DNA through its AT-hook-like motifs

To determine how BAZ1B can target to PCH, we expressed truncated BAZ1B mutants fused with EGFP in iMEFs (Fig. S4A). The C-terminal region containing a bromodomain (BrD) was found to be essential for BAZ1B localization to the PCH (Fig. 5A); however, the regions both upstream and downstream of the BrD, rather than the BrD as such, appeared to be critical for PCH targeting (Fig. S4A). Since neither H3K9me3 nor DNA methylation is required for PCH H3K36me2 (Fig. S2A and S2B), we hypothesized that BAZ1B directly binds to DNA elements. Then, we found that three AT-hook-like motifs, conserved across vertebrates (Fig. 5A), are located in the C-terminal region of BAZ1B (Fig. 5A; BrD+AT1-3). These motifs resemble the typical AT-hook motifs known to bind to AT-rich DNA (Arg-Gly-Arg-Pro; Aravind and Landsman, 1998). A BAZ1B mutant, which harbors amino acid substitutions in these AT-hook-like motifs (BrD+AT1-3mut) no longer localized to PCH (Fig. 5A). In addition, a truncated BAZ1B mutant (AT1-2) containing two AT-hook-like motifs but lacking BrD still targeted the PCH, although with a more diffuse background (Fig. S4A). These data suggest that three AT-hook-like motifs are required for DNA binding.

**Figure 5.**
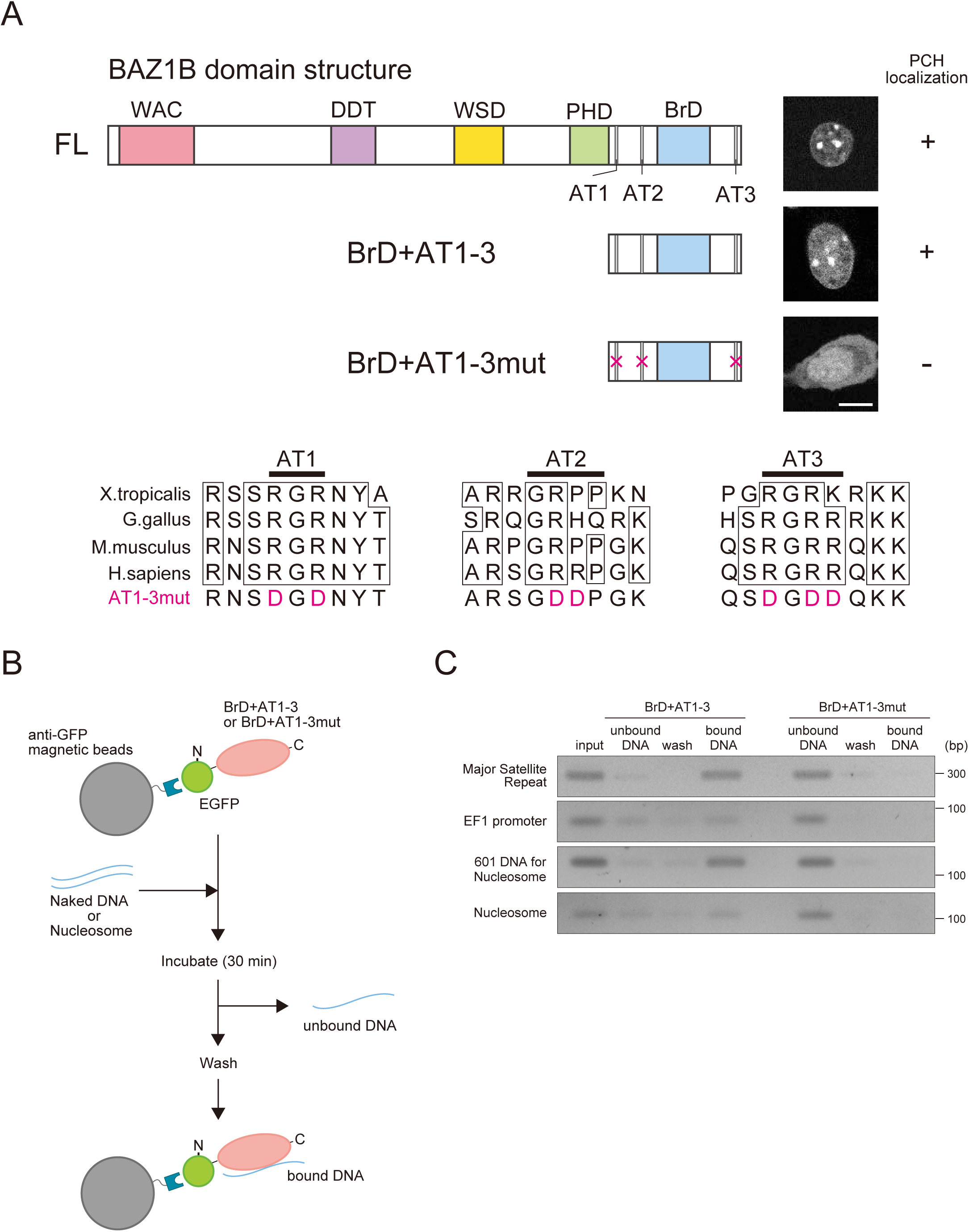
BAZ1B binds to AT-rich DNA through its AT-hook-like motif. (A) BAZ1B domain that targets PCH. Schematic drawing of the full length (FL), truncated BAZ1B consisting of bromodomain (BrD) and three AT-hook-like motifs (BrD+AT1-3), and those motifs mutant (BrD+AT1-3mut) is shown, with their representative live-cell images on the right. Amino acid sequence alignments of BAZ1B AT-hook-like motifs in different species and the substituted amino acids in AT1-3mut are shown. (B and C) DNA binding assay. (B) Experimental design. EGFP-tagged BAZ1B BrD+AT1-3 and BrD+AT1-3mut were expressed in HEK293T cells and purified using anti-GFP nanobody magnetic beads. After incubating BAZ1B domain-bound anti-GFP beads with naked DNA or nucleosomes for 30 min in 20 mM Tris-HCl [pH7.5] buffer, unbound DNA/nucleosomes were collected. After washing in 20 mM Tris-HCl [pH7.5] buffer containing 50 mM NaCl, bound-DNA/nucleosomes were eluted using 20 mM Tris-HCl [pH7.5] containing 350 mM NaCl. (C) Detection of DNA in different fractions by agarose-gel electrophoresis. The positions of size standards are indicated on the right. Scale bar, 10 μm.

We therefore examined whether the BAZ1B domain containing the three AT-hook-like motifs directly bind to AT-rich DNA in vitro (Fig. 5B). Both EGFP-BrD+AT1-3 and EGFP-BrD+AT1-3mut were expressed in HEK293T cells and then purified using anti-GFP magnetic beads (Fig. S4B). These beads were mixed with input DNA, either AT-rich major satellite repeats, GC-rich EF1α promoter, or 601 sequences (Lowary and Widom, 1998) (Fig. S4C-E). Following the removal of unbound DNA, the beads were washed, and the bound DNA was eluted (Fig. 5B). DNA from each fraction was then analyzed by agarose gel electrophoresis (Fig. 5C). Major satellite repeats bound to BrD+AT1-3 but not to BrD+AT1-3mut (Fig. 5C), whereas the binding of EF1α promoter to BrD+AT1-3 was much weaker. This result indicates that the AT-hook-like motifs have a preference for AT-rich DNA. We next examined whether BrD+AT1-3 could bind to nucleosomal DNA using a 145-bp 601 sequence. BrD+AT1-3 indeed bound to nucleosomes, although the binding affinity appeared to be weaker compared to the naked DNA (Fig. 5C). Thus, BAZ1B directly binds to AT-rich DNA via AT-hook-like motifs, which could recruit SMARCA5 and then NSD2 to induce H3K36me2 in AT-rich PCH.

### Coordinated expression of NSD2, SMARCA5, and BAZ1B promotes PCH localization of H3K36me2

To determine the molecular differences between cells with and without PCH H3K36me2, we investigated the protein expression levels of NSD2, SMARCA5, and BAZ1B by immunoblotting (Fig. 6A). Compared to cells without PCH H3K36me2 (i.e., MC12, C2C12, and Pmi28), NSD2 and SMARCA5 were highly expressed in NIH3T3 and iMEFs, both of which display high levels of H3K36me2 in PCH (Fig. 1C). BAZ1B was also highly expressed in iMEFs. In ESCs, SMARCA5 was relatively highly expressed, whereas levels of NSD2 and BAZ1b were moderate, possibly explaining the lower PCH enrichment of H3K36me2 in ESCs compared to iMEFs and NIH3T3 (Fig. 1C).

**Figure 6.**
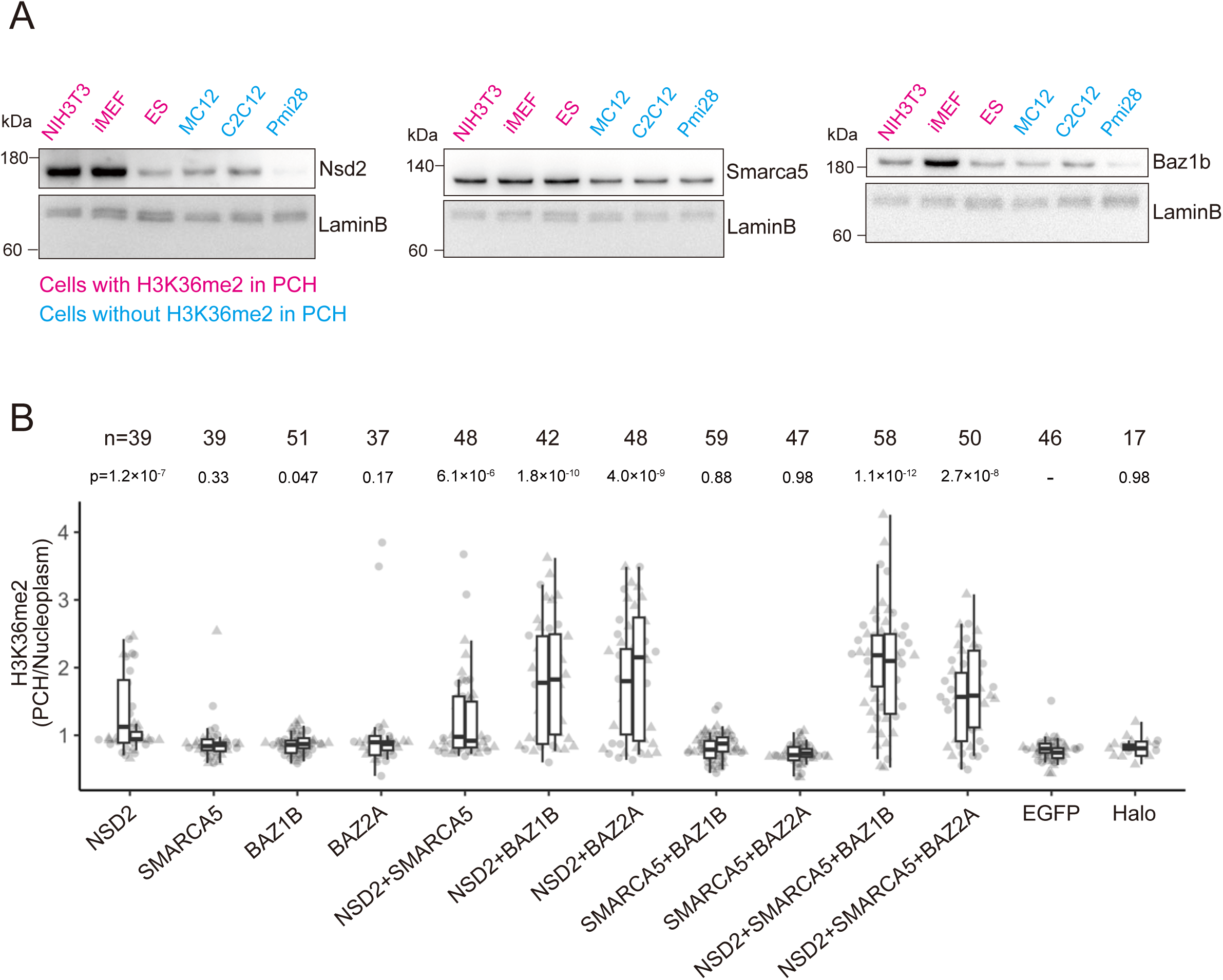
Cooperative high expression of NSD2, SMARCA5, and BAZ1B promotes PCH localization of H3K36me2. (A) Relative protein levels of NSD2, SMARCA5, and BAZ1B in mouse cell lines evaluated by Western blotting. Lamin B was detected as a loading control. The positions of size standards are indicated on the left. (B) PCH-to-nucleus intensity ratio of H3K36me2. Pmi28 cells were transfected with the expression vectors of indicated proteins, fixed, and stained with anti-H3K36me2, JF646 HaloTag ligand, and Hoechst33342. When expressing three proteins, an equal amount of each expression vector was mixed. When expressing one or two proteins, the same amount of each vector was used as was the case of triple expression, and the total DNA amount was adjusted using the EGFP- or Halo-expression vector. See Fig. 1 legend for the details of box plots. Note that p-values were calculated using the Mann-Whitney U test and Hommel correction with the EGFP control.

Using Pmi28 cells, which exhibit the lowest levels of NSD2 and BAZ1B without PCH H3K36me2, we investigated which proteins or their combinations, among NSD2, SMARCA5, and BAZ1B or BAZ2A, could induce PCH H3K36me2 by transient expression (Fig. 6B). When a single protein was expressed, only NSD2 induced a low level of PCH H3K36me2, compared to a control (EGFP alone), as shown before (Fig. 2D, 3E, 4D, and S3D). Combining SMARCA5 with either BAZ1B or BAZ2A did not have any effect. This result is reasonable because a methyltransferase is essential to increase H3K36me2. Co-expression of NSD2 with SMARCA5 did not markedly differ from the effects of NSD2 alone. By contrast, drastic increases of PCH H3K36me2 were observed when NSD2 was paired with either BAZ1B or BAZ2A. In addition, the concurrent expression of NSD2, SMARCA5, and BAZ1B further slightly increased PCH H3K36me2 levels. Since SMARCA5 levels are not very low in Pmi28, its overexpression appears to have a minimal impact on PCH H3K36me2 compared to the combined effects of NSD2 and BAZ1B, or BAZ2A. Taken together with KO analysis using iMEFs, these results indicate that PCH H3K36me2 is induced by the cooperation of NSD2, SMARCA5, and BAZ1B or BAZ2A.

### H3K36me2 is enriched in heterochromatin in mouse preimplantation embryos at the 2- to 4-cell stages

Finally, we investigated whether H3K36me2 is also localized to the PCH in mice. We first chose the spleen, lung, testis, liver, and brain for immunostaining, based on the high expression of at least two of NSD2, SMARCA5, BAZ1B, and BAZ2A, searching the public database (Palasca et al., 2018). However, none of these tissues showed H3K36me2 in PCH (Fig. S5A), suggesting that H3K36me2 is located in euchromatin in adult mouse tissues as generally observed. We also stained 13.5-day mouse embryo sections with anti-H3K36me2 but did not find PCH localization in any areas closely looked at (Fig. S5B).

We next analyzed totipotent cells in mouse preimplantation embryos since PCH H3K36me2 was found in undifferentiated ESCs. From the zygote to the 2-cell stage, the PCH is distributed in a ring around the nucleolus precursor body (NPB), and thereafter, the PCH gradually forms chromocenters until the blastocyst stage (Aguirre-Lavin et al., 2012). In paternal and maternal pronuclei (PN), H3K36me2 was not found in heterochromatin around NPBs, although H3K36me2 levels were higher in maternal PN (Fig. 7A and S5C). At both the 2- and 4-cell stages, however, H3K36me2 was in part concentrated in heterochromatin around NPBs and chromocenters (Fig. 7A and S5C). Indeed, colocalization analysis of individual nuclei revealed that H3K36me2 at the 2- and 4-cell stages exhibited a higher correlation with DNA compared to other stages (Fig. 7B). Although H3K36me2 was not concentrated on PCH in most cells during the morula and the blastocyst stages, a small number of nuclei in blastocyst inner cell mass exhibited some enrichments in PCH (Fig. 7A and 7B). These results indicate that H3K36me2 can be enriched in heterochromatin at limited developmental stages in vivo. As the heterochromatin in mammalian preimplantation embryos is transcribed and dynamically reorganized, the deposition of H3K36me2 might be involved in these processes.

**Figure 7.**
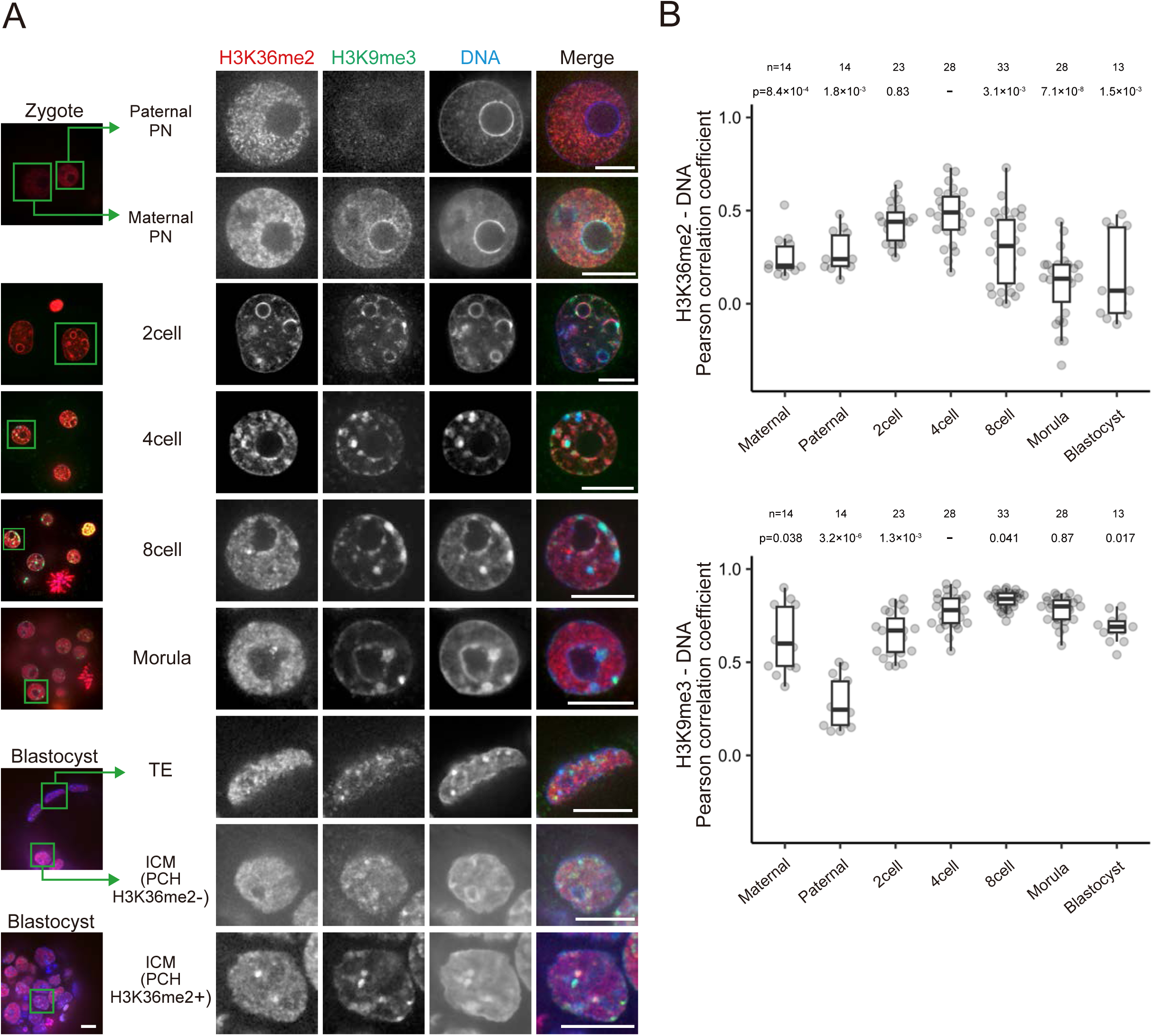
H3K36me2 is distributed in a ring-shaped PCH around the NPB in mouse preimplantation embryos. Localization of H3K36me2 in mouse preimplantation embryos at various stages. Embryos were fixed and stained with anti-H3K36me2 (red), anti-H3K9me3 (green), and Hoechst 33342 (blue). (A) Images of single optical sections. Low-power views of merged images are shown on the left. Magnified views of indicated nuclei are shown on the right. PN, pronucleus; TE, trophectoderm; and ICM, inner cell mass. (B) Pearson correlation coefficients of DNA and H3K36me2 (top) or H3K9me3 (bottom). The number of nuclei (n) and p-values, calculated using the Mann-Whitney U test and Hommel correction to the 4-cell stage, are shown. See Fig. 1 legend for the details of box plots. Scale bars, 10 μm

## Discussion

Here we showed that H3K36me2, typically associated with euchromatin, is enriched in heterochromatin in some mouse cell lines and preimplantation embryos at the 2- to 4-cell stages. Overexpression and KO experiments revealed that H3K36me2 localization at PCH is regulated through NSD2, SMARCA5, and BAZ1B or BAZ2A.

### Mechanisms guiding H3K36me2 to heterochromatin

PCH histone modifications, such as H4K20me3, typically occur in an H3K9me3-dependent manner (Schotta et al., 2004). The PCH localization of H3K36me2 was, however, independent of either H3K9me2/3 or DNA methylation. We found that NSD2, the enzyme responsible for H3K36me2, targets PCH through the ISWI complexes that harbor subunits with AT-rich DNA binding motifs. We showed that BAZ1B, which forms the WICH with SMARCA5, has three AT-hook-like motifs intervened by a BrD at its C-terminus. Although these motifs are not typical AT-hook, they are essential for DNA binding, in agreement with previous studies showing that variations in the AT-hook core and surrounding amino acids are somewhat permissive (Aravind and Landsman, 1998; Filarsky et al., 2015). Additionally, BAZ2A, which forms the NoRC with SMARCA5, was also localized to heterochromatin and assists in PCH H3K36me2 localization. BAZ2A has been reported to bind to DNA through its Tip5/ARBP/MBD (TAM) domain with AT-hook motifs (Chen et al., 2021; Filarsky et al., 2015). Hence, these ISWI complexes may redundantly function in recruiting NSD2 for PCH H3K36me2 localization. Even though BAZ1A could target PCH via its N-terminal WAC domain (Tate et al., 1998), we did not observe PCH localization of emGFP-BAZ1A.

In Pmi28 cells, which display euchromatic H3K36me2 localization and express relatively low levels of BAZ1B and NSD2, the ectopically expressed emGFP-BAZ1B localized to PCH perhaps via its AT-hook-like motifs that bind to DNA. However, as BAZ family proteins possess various chromatin binding domains such as PHD and BrD, their binding affinity and specificity to DNA and histones could be modulated by chromatin modification context (Tallant et al., 2015; Oppikofer et al., 2017), which may in turn result in diverse localization in different cell types. The efficiency of ISWI complexes targeting PCH may also depend on the cell cycle since both BAZ1B and SMARCA5 exhibit heterochromatin localization from mid to late S phase (Collins et al., 2002; Bozhenok et al., 2002; Poot et al., 2004). The association of BAZ1B with newly replicated chromatin could be mediated through specific histone modifications and/or increased DNA accessibility during replication since the AT-hook-like motifs bind to naked DNA more strongly than to nucleosomes.

Of note, a recent report suggests that the histone methyltransferase DOT1L introduces H3K79me3 to PCH in a SMARCA5-dependent manner in mouse ESCs (Malla et al., 2023). Although the SMARCA5’s partner in this process remains unidentified, the ISWI complexes might decorate PCH by attracting various modifying enzymes.

### Physiological relevance of H3K36me2 on heterochromatin

The relatively higher expression of NSD2, SMARCA5, and BAZ1B appears to be associated with H3K36me2 PCH localization in mouse cell lines, but there was no obvious common feature among cells showing PCH H3K36me2; for example, whereas PCH H3K36me2 in ESCs suggests its relevance to pluripotency, differentiated fibroblasts also showed the same localization. To explore the physiological relevance of PCH H3K36me2, we examined adult mouse tissues and embryos. So far, heterochromatic H3K36me2 was exclusively observed in preimplantation embryos, particularly at the 2- and 4-cell stages. A minor fraction of cells in blastocysts also exhibited PCH H3K36me2 might be related to ESCs. In mice, the major zygotic genome activation undergoes at the 2-cell stage, including transient transcription from major satellite repeat (Probst et al., 2010), rRNA (Yu et al., 2021), retrotransposons LINE (Jachowicz et al., 2017), and MERVL (Sakashita et al., 2023). Appropriate transcription from these repeats is important for normal development. PCH H3K36me2 at these stages may fine-tune heterochromatin transcription as previously suggested (Frescas et al., 2008; Suzuki et al., 2016; Chaouch et al., 2021). H3K36me2 may also play a role in heterochromatin reorganization in embryogenesis mediated through the perinucleolar ring-shaped structures at the 2-cell stage to more typical chromocenters at the later stages. A very recent study has shown a model that H3K9me3 and H3K36 methylation tandemly promote DNA methylation (Sinha et al., 2023). DNA methylation in oocytes is also regulated by H3K36me2 and H3K36me3 (Yano et al., 2022). As both H3K9me3 and DNA methylation are dynamically changed during these stages, H3K36me2, which is targeted to heterochromatin independently of H3K9me3, might play a critical role in DNA methylation on heterochromatin repeat sequences. To support this notion, H3K36me2 localization in ring-shaped heterochromatin was also observed in porcine 2-cell stage embryos, although the zygotic genome activation occurs at the later stages (Diao et al., 2014). H3K36me2/3 is also known to cross-talk with other chromatin modifications, such as antagonistically distributing H3K27me3 (Streubel et al., 2018; Chen et al., 2022), during oocyte maturation and zygotes (Xu et al., Nat Genet, 2019). Transient accumulation of H3K36me2 might prevent H3K27me3 invasion into constitutive heterochromatin before H3K9me3 distribution is established. Further analysis will be required to elucidate the role of H3K36me2 in embryonic development.

## Materials and Methods

### Plasmid construction

The following Kazusa HaloTag-tagged human cDNA expression vectors were used: NSD2 (FHC00177), NSD3 (FHC00329), SETD2 (FHC00272), SETD3 (FHC12875), SETD7 (FHC00270), SETD8 (FHC06789), ASH2L (FHC01483), SMYD2 (FHC27765), SMYD3 (FHC03496), SETMAR (FHC05461), KDM2A (FHC 00712), KDM3C (FHC36094E), KDM4A (FHC00602), KDM4B (FHC13945E), KDM4C (FHC00635), and SMARCA5 (FHC01216). For expressing human BAZ1A, BAZ1B, and BAZ2A, we used the following emerald-green GFP (emGFP)-tagged vectors from Addgene: BAZ1A (#65371), BAZ1B (#65372), and BAZ2A (#65373). To construct the super-folder GFP (sfGFP)-tagged SMARCA5 expression vector, the ORF was amplified by PCR using Halo-SMARCA5 (#FHC01216; Kazusa) as a template and inserted into sfGFP-N1 vector (#54737; Addgene) digested with EcoRI using the In-Fusion HD Cloning Kit (Takara Bio). Then, SMARCA5-sfGFP was cloned into the PB533A-2 vector (Systems Biosciences). To identify the BAZ1B domain required for PCH localization, enhanced GFP (EGFP)-tagged truncated BAZ1B expression vectors were constructed. The full-length and domain of interest were amplified by PCR using emGFP-BAZ1B (#65372; Addgene) as a template and inserted into pEGFP-C2 vector (Takara Bio) digested with EcoRI using the In-Fusion HD Cloning Kit. One truncated vector without the PHD finger and bromodomain (BrD) (emGFP-ΔPHD, BrD) was produced by inverse PCR using emGFP-BAZ1B as a template. To generate a truncated vector with BrD and three mutated AT-hook-like motifs (EGFP-BrD+AT1-3mut), site-directed mutagenesis was performed by inverse PCR using EGFP-BrD+AT1-3 as a template. The primers used are listed in Table S1.

### Animal care

Mouse care and experimental procedures were approved by the Institutional Animal Experiment Committee of the Tokyo Institute of Technology (Application number: D2023004) and the Animal Care and Use Committee of Kindai University (Application number: KAAT-22-001). All animal experiments were performed in accordance with institutional and governmental guidelines. Mice were maintained on a 12:12 dark: light cycle at a constant temperature of 22–23°C and were fed freely with food and water. For preparing adult mouse tissue specimens, we used 26-month-old C57BL/6 male mice that were housed in ventilated cages and kept in a pathogen-free, temperature-controlled facility at the Tokyo Institute of Technology.

### Cell culture

NIH3T3 mouse embryonic fibroblasts (obtained from N. Takagi at Hokkaido University), MC12 mouse embryonic carcinoma cells (obtained from N. Takagi at Hokkaido University; Abe et al., 1988), C2C12 myoblasts (obtained from JCRB), and HEK293T cells (obtained from K. Fujinaga at Sapporo Medical School) were grown in high-glucose Dulbecco’s modified Eagle’s medium (DMEM; Nacalai Tesque) with supplements (1% penicillin/streptomycin [Nacalai Tesque] and 10% fetal bovine serum [FBS; Thermo Fisher Scientific, Gibco]). Immortalized mouse embryonic fibroblasts (iMEFs; obtained from Y. Shinkai at RIKEN), H3K9me-related enzymes knockout (5KO) iMEFs (obtained from Y. Shinkai; Fukuda et al., 2023) were grown in high-glucose DMEM with supplements (1% penicillin/streptomycin, 10% FBS, 1% Non-essential Amino Acids [NEAA; Wako], 55 µM 2-Mercaptoethanol [Gibco]). Mouse embryonic stem (ES) cells (TT2; obtained from Y. Shinkai) were grown in high-glucose DMEM with supplements (1% penicillin/streptomycin, 10% FBS, 1% NEAA, 1% L-glutamine [Gibco], 10^3^ units/ml leukemia inhibitory factor [Nacalai Tesque], 100 µM 2-Mercaptoethanol). Triple DNA methyltransferase knockout (TKO) ES cells (obtained from M. Okano; Tsumura et al., 2006) were grown in high-glucose DMEM with supplements (1% penicillin/streptomycin, 15% FBS, 1% NEAA, 1 mM sodium pyruvate [Gibco], 10^3^ units/ml leukemia inhibitory factor, 100 µM 2-Mercaptoethanol). Pmi28 myoblasts were grown in Ham’s F-10 medium with supplements (1% penicillin/streptomycin, 20% FBS). All of the cells were grown at 37°C under a 5% CO_2_ atmosphere.

### Generation of Knockout (KO) cells

The CRISPR/Cas9 system was used for KO cell establishment. All guide RNAs (gRNAs) were designed by CRISPRdirect (https://crispr.dbcls.jp/). The DNA sequences for gRNAs are listed in Table S1. Annealed oligonucleotides were inserted into pSpCas9(BB)-2A-Puro (PX459) V2.0 vector (#62988; Addgene) digested with BbsI. iMEFs were transfected with gRNA vectors using Lipofectamine 3000 (Thermo Fisher Scientific). For the NSD2 knockout, two gRNAs targeting the PWWP and SET domains were used. The next and 3 days later, the medium was replaced with the one containing 2 µg/mL puromycin. For single-cell cloning, puromycin-selected cells were seeded at a density of 500 cells/20 mL in a 15-cm dish to form colonies. Then, single colonies were picked and transferred to a glass-bottom (for immunofluorescence) or plastic-bottom (for stock) 96-well plate. To screen KO-candidate cells for each gene, immunostaining was performed using the following antibodies: anti-H3K36me2 for NSD2-KO, anti-SMARCA5 for SMARCA5-KO, and anti-BAZ1B for BAZ1B-KO cells. The depletion of each protein in KO cells was then validated by Western blotting.

### Generation of Knockdown (KD) cells

The shRNA sequences were designed using the GPP Web Portal (https://portals.broadinstitute.org/gpp/public/). shRNA sequences are listed in Table S1. Two shRNAs were used for each gene knockdown. Annealed oligonucleotides were inserted into the pLKO.1-Hygro vector (#24150, Addgene) digested with AgeI and EcoRI. The day after seeding HEK293T cells on 6-cm dishes, pLKO.1-Hygro-based vector, psPAX2 (#12260, Addgene), and pVSV-G (#PT3343-5, Clontech) vectors were transfected using Lipofectamine 3000. The following day, the medium was replaced with fresh medium, which was then collected 48 h post-transfection. Then, iMEFs were incubated with culture supernatant containing the virus and 4 μg/mL polybrene (Nacalai Tesque) for 24 h. Infected cells were selected in a medium containing 400 μg/mL hygromycin B Gold (InvivoGen) for 3 days.

### Transient transfection and live-cell imaging

For Fig. 2C, S2C, 3C, 3D, 3I, 4H, and 4I, cells were plated on a glass bottom 24-well plate (AGC Techno Glass). The following day, HaloTag-, sfGFP-, emGFP-, or EGFP-tagged expression vectors were transfected using Lipofectamine 3000. After 24 h of regrowth, cells were incubated in a medium containing 2 nM Janelia Fluor 646 (JF646) HaloTag ligand (Promega) and 0.5 μg/mL Hoechst33342 (Nacalai Tesque) for 30 min. For Fig. S3C, 4C, S4A, and 5A, cells were only stained with Hoechst33342. After washing twice with FluoroBrite medium (Thermo Fisher Scientific) containing the supplements, the plate was positioned on a heated stage (Tokai Hit) at 37°C under 5% CO_2_ on a point-scan confocal microscope (Ti2 with A1 system; Nikon) operated by the built-in software NIS-elements AR version 5.21.00 with a Plan Apo VC 60x WI DIC N2 (NA 1.2) objective, water immersion lens (Nikon). Images were acquired using a 405-nm, 488-nm and a 640-nm laser lines (LU-N4; Nikon), GAsP detectors, a 405/488/561/640 dichroic mirror, and 450/50, 525/50, 595/50, 700/75 emission filters, with the following parameters; Z range, 5 µm; Z step, 1 µm; Pinhole size, 43.42 µm; Scanner zoom, 1.5x; Scan size, 512 x 512 pixels; and Scan speed, 1 frame/sec.

### Immunofluorescence for cultured cells

For immunostaining of histone marks, cells were fixed with 4% formaldehyde in 250 mM HEPES–NaOH (pH 7.4) and 0.1% Triton X-100 for 5 min at room temperature. After washing with PBS, cells were permeabilized with 1% Triton X-100 in PBS for 20 min at room temperature. After blocking with Blocking One-P (Nacalai Tesque) for 20 min at room temperature, cells were incubated with 2 μg/mL mouse monoclonal antibody directly conjugated with a fluorescence dye (Alexa Fluor 488, Cy3, or Cy5) and 1 μg/mL Hoechst33342 overnight at 4°C. After washing with PBS three times, fluorescence images were collected. Antibodies used are listed in Table S2. For Fig. S1B, cells were stained with 2 μg/mL rabbit anti-H3K36me2 monoclonal antibody, Cy3-conjugated mouse anti-H3K9me3 monoclonal antibody, and 1 μg/mL Hoechst33342 overnight at 4°C, and then incubated with 1 μg/mL Alexa Fluor 488-conjugated anti-rabbit antibody for 1 h at room temperature.

For Fig. 2A, 2B, 2D, 2F, 2G, 3E, 3G, and 3H, 24 h after transfection with HaloTag expression vectors, cells were fixed with 4% formaldehyde in PBS for 5 min at room temperature. After washing with PBS, cells were permeabilized with 1% Triton X-100 in PBS for 20 min at room temperature. After blocking with Blocking One-P (Nacalai Tesque) for 20 min at room temperature, cells were incubated with 2 μg/mL fluorescence dye-conjugated mouse monoclonal antibody, 2 nM JF646 HaloTag ligand, and 0.5 μg/mL Hoechst33342 overnight at 4°C. For Fig. 4B, 4D, 4F, 4G, S3A, S3B, and S3D, cells were stained without the HaloTag ligand 24 h after transfection with emGFP-tag expression vectors. For Fig. 6B, up to three different plasmids were mixed; in the case of using only one or two expression vectors, the total amount of DNA was equalized using EGFP or Halo vectors. Cells were fixed 48 h after transfection for immunostaining.

For Fig. 1A, 1B, S1C-E, fluorescence images were collected using a high-resolution spinning-disk confocal microscope (Ixplore SpinSR; Olympus) operated by the built-in software CellSens Dimension 3.2, equipped with a UplanApoN 60× OSC2 (NA 1.4) objective, oil immersion lens at room temperature. Images were acquired using 405-, 488-, 561-, and 640-nm laser lines (OBIS), a 405/488/561/640 dichroic mirror, and 447/60, 525/50, 617/73, and 685/40 emission filters, a scientific complementary metal oxide semiconductor (sCMOS) camera (ORCA Flash 4.0; Hamamatsu Photonics) with the following parameters; Z range, 8 µm; Z step, 0.5 µm; Scan size, 2048 x 2048 pixels; Camera adapter magnification, 1x.

For other figures, fluorescence images were collected using a point-scan confocal microscope (Ti2 with A1 system; Nikon) operated by the built-in software NIS-elements AR with a Plan Apo VC 60x WI DIC N2 (NA 1.2), water immersion objective lens at room temperature. Images were acquired using 405-, 488-, 561-, and 640-nm laser lines (LU-N4; Nikon). Other setting parameters were the same as described in "Transient transfection and live-cell imaging.”

### Immunofluorescence for mouse tissue and E13.5 embryo sections

Mice were euthanized by cervical dislocation. Tissues of interest and E13.5 embryos were isolated and washed with PBS. Tissues and embryos were fixed in 4% formaldehyde in PBS overnight at 4°C. After washing with PBS, tissues were soaked in the optimal cutting temperature compound (SAKURA) and frozen in liquid nitrogen. Frozen tissues were sectioned at 10 µm thick using a microtome (CM3050S; Leica). The sections were attached to glass slides, washed with PBS-T (0.05% Tween20 in PBS), and heated at 95°C for 20 min in 10mM citric acid (pH2.0) for antigen retrieval using a decloaking chamber NxGen (FUNAKOSHI). After washing with PBS-T, the sections were permeabilized with 1% Triton X-100 in PBS for 20 min, washed with PBS-T, and incubated with Blocking One-P for 20 min at room temperature. Then, the sections were stained with anti-H3K36me2 Fab conjugated with Alexa Fluor 488 (20 μg/mL) and the anti-H3K9me3 antibody conjugated with Cy3 (for adult mouse sections) or with Cy5 (for embryo sections) (10 μg/mL), and 1 μg/mL Hoechst33342 for three days at room temperature. After washing with PBS-T, the samples were mounted with Prolong Diamond (Thermo Fisher Scientific) and covered with a coverslip (MICRO COVER GLASS 24×32mm, MATSUNAMI).

Fluorescence images for adult tissue sections were acquired using a point-scan confocal microscope (Ti2 with A1 system; Nikon) with a Plan Apo VC 60x WI DIC N2 (NA 1.2) objective lens as described above except Zoom factor (1x). Images were acquired using 405-488-, 561-, and 640-nm laser lines (LU-N4; Nikon). Other setting parameters were the same as described in "Transient transfection and live-cell imaging.”

Fluorescence images for embryo sections were acquired using a spinning disk confocal microscope (CSU-W1; Yokogawa) operated by the built-in software NIS-elements v5.11.03, equipped with an inverted microscope (Ti-E; Nikon), EM-CCD (iXon 3; Andor), a laser illumination system (LDI-NIR; Chroma Technology Japan), and a Plan Apo 40x/0.95 DIC M/N2 dry objective lens at room temperature. Images were acquired using 405-, 470-, and 640-nm laser lines, a 405/470/555/640 dichroic mirror, and 440/40, 520/60, and 690/50 emission filters, with the following parameters; Z range, 5 µm; Z step, 1 µm; Scan size, 1024 x 1024 pixels.

### In vitro fertilization and immunofluorescence of mouse preimplantation embryos

For preparing preimplantation embryos, mice were euthanized with cervical dislocation immediately before oocyte and sperm collection. Cumulus-intact oocytes were collected in 0.2 mL of TYH medium and inseminated with capacitated sperm at a final concentration of 75 sperm/μL. After 1.5 h incubation at 37 °C under 5% CO_2_ in the air, cumulus cells were dispersed by a brief treatment with hyaluronidase (Type-IS, 150 units/mL, Sigma). The denuded fertilized oocytes were transferred to KSOMaa medium and incubated until they reached the appropriate stages.

The embryos were fixed with 3.7% paraformaldehyde containing 0.2% Triton X-100 for 20 min at room temperature. After fixation, the embryos were washed twice with PBS containing 3% bovine serum albumin (BSA) and placed for 60 min at room temperature for blocking. The embryos were then incubated with anti-H3K9me3 antibody conjugated with Alexa Fluor 488 (10 μg/mL) and anti-H3K36me2 antibody conjugated with Cy3 (5 μg/mL) dissolved in the above blocking buffer at 4 °C for at least 8 h. After washing with PBS, the embryos were transferred to PBS containing 0.5 μg/mL of DAPI (Nacalai Tesque) in a glass-bottomed dish (Mat Tek, P35G-1.5-14-C).

The fluorescent observations were performed using a spinning disk based-super resolution confocal microscopic system (CSU-W1 SoRa; Yokogawa Electric Corp.; Hatano et al., 2022) equipped with an inverted microscopy (IX73; Olympus) with a 40× silicone immersion objective lens (UPLSAPO40XS: NA 1.25; Olympus) and an intermediate 4× tube lens. Three-color fluorescence images in 100 different focal planes at 0.5 μm intervals were captured with 405-, 488-, and 561-nm laser lines. The images were passed through emission filters (FF02-460/80 for DAPI, FF01-520/35 for Alexa488, and FF02-617/73 for Cy3; Yokogawa) and captured by a Prime95B sCMOS camera (Teledyne Photometrics) with de-noised switch. The system was controlled and images were taken by μ-Manager microscopy software (Edelstein et al., 2014).

### Image analysis

Line profiles were drawn using NIS-Elements AR Analysis version 5.01.00 (Nikon) and Image J Fiji version 1.54f (https://fiji.sc/). A merged image from single Z-slice images of different fluorescence channels was created using NIS-Elements. Using the "Plot Profile" function in Fiji, a straight line was drawn on the image, allowing for the measurement of both position and intensities. A line profile plot was then generated using R version 4.2.1 (https://www.r-project.org/), with the x-axis representing the distance (µm) and the y-axis representing the intensity.

Intensity measurements were performed using NIS-Elements (Nikon) and Aivia version 10.5.1 (DRVision). After subtracting the background with NIS-Elements, the Aivia "Cell Analysis" recipe was run to define regions for chromocenters and cell nuclei based on Hoechst signals, and then mean pixel intensities were measured. Fold enrichment in PCH (the mean intensity in the chromocenter region divided by that of the nucleus excluding the chromocenter) was calculated using R.

Pearson correlation coefficients between two fluorescence intensities in single nuclei were calculated using the Image J Fiji “Coloc2” function. Plots and significance tests were performed using R.

### Quantitative RT-PCR (RT-qPCR)

Total RNA was isolated from cells using TRIzol (Thermo Fisher Scientific) and further purified using an RNeasy MinElute Cleanup Kit (Qiagen) according to the manufacturer’s instructions. cDNA was synthesized by reverse transcription using Superscript III Reverse Transcriptase (Thermo Fisher Scientific) and random hexamer primers. cDNA was mixed with PowerUP SYBR Green Master Mix (Applied BioSystems) and forward and reverse primers (Table S1) in 96-well format tubes, before setting to a real-time PCR system (Agilent) with the following parameters: Initial hold, 50°C for 2 min and 95°C for 2 min; PCR, (95°C for 15 sec and 60°C for 60 sec) x 40 cycles; Dissociation curve, 95°C for 60 sec, 55°C for 30 sec, and 95°C for 30 sec. For relative quantification, standard curves were prepared for the target genes and the reference gene (Gapdh) using cDNA from NIH3T3 cells. The initial template amounts of the target and reference genes were calculated based on the Ct values, and the expression levels of the target genes were relative to the reference gene. Two reaction tubes were prepared for each of the two biological replicates.

### Protein extraction and immunoprecipitation

HEK293T cells grown on two 10-cm dishes were transfected with an expression vector for EGFP-BrD+AT1-3 or EGFP-BrD+AT1-3mut using Lipofectamine 2000 (Thermo Fisher Scientific). The next day, cells were washed with PBS, trypsinized, and collected in a 15-mL tube by centrifugation (1,200 rpm for 2 min). The cell pellet was suspended in ice-cold PBS, transferred to a 1.5 mL tube, and washed with ice-cold PBS three times, before lysis in Lysis buffer (300 mM NaCl, 10 mM HEPES-NaOH [pH7.4], 0.1% TritonX-100, 1× Protease inhibitor cocktail [Nacalai Tesque]). After centrifugation (20,000 x g for 20 min at 4°C), the supernatant was transferred to a new 1.5 mL tube. GFP-Trap Magnetic Agarose beads (Proteintech; 80 µL) were prewashed with wash buffer (10 mM Tris-HCl [ph7.5], 150 mM NaCl, 0.5 mM EDTA) in a 1.5 mL tube. After collecting beads using a magnetic rack (Thermo Fisher Scientific), the wash buffer was removed. This washing step was repeated a total of three times. The cell lysate was then added to the GFP-Trap beads and incubated at 4°C overnight with gentle mixing. After washing the beads three times with wash buffer, they were resuspended in the same buffer. Small aliquots were collected from the input cell lysate, washed supernatant, and bead suspension for SDS-polyacrylamide gel electrophoresis to evaluate purity and quantity. Aliquots were mixed with an equal volume of 2×SDS-gel loading buffer (4% SDS, 20% glycerol, 120 mM Tris-HCl [pH 6.8], 100 mM DTT, 0.01% bromophenol blue) and heated at 95°C for 5 min. Proteins were then separated on a 15% polyacrylamide gel (Supersep Ace, Fujifilm) alongside a dilution series of BSA for quantitation. The concentrations of the purified EGFP-BrD+AT1-3 or EGFP-BrD+AT1-3mut proteins were measured using a standard curve based on known amounts of BSA.

### DNA-binding assay

AT-rich Major Satellite repeats were amplified by PCR using specific primers (Table S2) and NIH3T3-derived DNA as a template. PCR products were electrophoresed on an agarose gel, and ∼250 bp fragments were gel extracted for cloning using a PCR Cloning Kit (NEB). The resulting plasmid was purified by Midiprep (Thermo Fisher Scientific), and the insert sequence was determined by Sanger sequencing. After EcoRI digestion and agarose gel electrophoresis, the insert Major Satellite repeats (289 bp) were gel extracted and purified using a Gel-extraction kit (Qiagen). For preparing GC-rich DNA fragments, the EF1α promoter flanking sequence (78 bp, 67% GC contents) from the PB533A-Halo vector was utilized by AsiSI and MfeI digestion and gel extraction. The preparation of Widom 601 DNA and nucleosome was described previously (Arimura et al., 2012; Vasudevan et al., 2010; Kujirai et al., 2018). Nucleotide sequences used for binding assay are shown in Fig. S4C-E.

For the DNA binding assay, GFP-Trap Magnetic Agarose beads that captured EGFP-BrD+AT1-3 and EGFP-BrD+AT1-3mut proteins were prewashed with High-salt buffer (1 M NaCl, 20 mM Tris-HCl [pH 7.5], 1 mM DTT) to eliminate any residual genomic DNA, before washing twice with Reaction buffer (20 mM Tris-HCl [pH7.5], 1 mM DTT). After removing the Reaction buffer, DNA or nucleosomes (0.28 pmol) in 10 µL of Reaction buffer were added to the beads (0.14 pmol), and the mixture was incubated for 30 min at room temperature with gentle rotation. The tubes were placed on a magnetic rack, and the supernatant was collected. Beads were washed with Wash buffer (50 mM NaCl, 20 mM Tris-HCl [pH7.5], 1 mM DTT) for 5 min at room temperature before the supernatant was collected. Then, Elution buffer (350 mM NaCl, 20 mM Tris-HCl [pH 7.5], 1 mM DTT) was added, incubated at room temperature for 5 min, and the supernatant was collected. All supernatants were electrophoresed on a 2% agarose gel and stained with Gel Red (Biotium).

### Western blotting

The day after cells were plated on a 6-cm dish, cells were washed three times with ice-cold PBS and 500 µL of 2×Laemmli buffer (4% SDS, 20% glycerol, 120 mM Tris-HCl [pH 6.8]) was added. Cell lysate was harvested using a cell lifter (Corning) and transferred to a 1.5 mL tube. After mixing with DTT (0.1 M) and bromophenol blue (0.01%), tubes were heated at 95°C for 10 min, proteins were separated in 7.5% SDS-polyacrylamide gels (SuperSep™ Ace; Fujifilm) and transferred to polyvinylidene fluoride membranes (FluoroTrans W; Pall) with EzFastBlot transfer buffer (ATTO) using a transfer system (Trans-Blot Turbo; Bio-Rad). After washing with TBS-T (10 mM Tris-HCl [pH 8.0], 0.1% Tween-20, 150 mM NaCl), membranes were incubated with Blocking One (Nacalai Tesque) for 20 min and then incubated with primary antibodies diluted in Can-Get-Signal Solution I (TOYOBO) for 2 h at room temperature. After washing in TBS-T for 10 min three times, membranes were incubated with horse radish peroxidase-labeled secondary antibody (Jackson ImmunoResearch) diluted in Can-Get-Signal Solution II (TOYOBO) for 1 h at room temperature. After washing membranes with TBS-T for 10 min three times, chemiluminescence signals were developed using ImmunoStar LD (Wako) and detected using a LuminoGraph II Chemiluminescent Imaging System (ATTO). Antibodies used are listed in Table S2.

### ChIP-seq data analysis

ChIP-seq dataset of H3K36me2 and H3K9me3 were obtained from the National Center for Biotechnology Information Sequence Read Archive (NCBI SRA) for ESCs (PRJNA720715), B cells (PRJNA324130), and pancreatic cancer cells (PRJNA565773). The fastq data were preprocessed with fastp ver. 0.23.2 and mapped onto the mouse mm39 genome using bowtie2 ver. 2.4.5 with the "very-sensitive-local" option. After removing unmapped read data using sambamba ver. 0.6.6, duplicated reads were removed using the markdup function in samtools ver. 1.13. The mapped data were utilized to generate bigwig files using bamCoverage function of deepTools ver. 3.2.1 with binSize of 20 and RPKM normalization. The correlations among the ChIP-seq data were calculated for consecutive genome bins using multiBigwigSummary bins function with 1,000 binSize, and the Spearman correlation was visualized with deepTools plotCorrelation.

Regarding transposable elements, RepeatMasker output for the mouse mm39 genome (accessed on May 14, 2023) was obtained from the UCSC Genome Browser. For each class of SINE, LINE, LTR retrotransposons, and DNA transposons, relatively young copies (<10% divergence from consensus) that were nearly full-length (>90% in length of consensus) were selected for subsequent analysis. The deepTools computeMatrix tool was used to calculate the accumulation of ChIP-seq signals within ±3 kbp regions of transcription start sites (TSS) or the TSS-TES (transcription end sites) of the transposable elements. The resulting data were visualized using the deepTools plotHeatmap tool.

### Amino acid alignments

BAZ1B amino acid sequences of H. sapiens (NP_115784.1), M. musculus (NP_035844.2), G. gallus (XP_001233717.3), X. tropicalis (XP_004911823.1), and D. rerio (XP_003200529.2) were aligned using GENETYX version 11 software (Genetyx).

### Statistical analysis

Statistical significance tests were performed using R. For comparisons between two groups, the Mann-Whitney U test was applied. For comparisons of three or more groups, the Mann-Whitney U test with Hommel correction was used.

### Artificial Intelligence

For English proofreading, Grammarly and ChatGPT-4 (Open AI), with a prompt “The following document is a part of a scientific paper. Please proofread and correct any grammatical errors.”, were in part used. No original sentences were generated using AI chatbots.

### Online supplemental material

Fig. S1 shows H3K36me2 distribution in various mouse cells, related to Fig. 1. Fig. S2 shows the results of screening of factors that are involved in PCH H3K36me2 localization, including the localization of H3K36me2 in cells lacking H3K9me3 and DNA methylation and the localization of Halo-tagged methyltransferases and demethylase, related to Fig. 2. Fig. S3 shows the localization of BAZ1A and BAZ2A, related to Fig. 4. Fig. S4 shows the details for BAZ1B-DNA interaction, including the localization of various truncated proteins, purification of the DNA binding domain, and DNA sequences used for the binding assay, related to Fig. 5. Fig. S5 shows immunofluorescence data for mouse adult tissue and embryo sections, and line plots for zygote and 2-cell embryos. Tables S1 and S2 list oligonucleotides and antibodies with the conditions used in this study, respectively.

## Acknowledgments

We thank Y. Shinkai, M. Okano, N. Takagi, and K. Fujinaga for providing cell lines and members of the Kimura and Yamagata Labs for help and discussion, especially H. Oda for assisting with an imaging analysis and Y. Kono and T. Shimi for instructing the lentivirus system. We also thank the Center for Integrative Biosciences and the Biomaterials Analysis Division, Open Facility Center at Tokyo Institute of Technology for DNA sequencing.

This work was supported by Japan Society for the Promotion of Science KAKENHI (JP17H01417 and JP21H04764 to H. Kimura, JP18H05528 to K. Yamagata, JP20H05690 to T. Kujirai, JP23H05475 to H. Kurumizaka, and JP22K06338 to H. Nishihara), Japan Science and Technology Agency CREST (JPMJCR20S6 to Y. Sato and JPMJCR16G1 to H. Kimura) and ERATO (JPMJER1901 to H. Kurumizaka), and Japan Agency for Medical Research and Development (AMED) Basis for Supporting Innovative Drug Discovery and Life Science Research (BINDS) (JP22ama121020 to H. Kimura and JP23ama121009 to H. Kurumizaka).

The authors declare no competing financial interests.

## Author contributions

Conceptualization, N. Goto, H. Kimura; investigation, N. Goto, K. Suke, N. Yonezawa, H. Nishihara; formal analysis, N. Goto; methodology, Y. Sato; resources, T. Kujirai, H. Kurumizaka; writing – original draft, N. Goto; writing – review and editing, K. Suke, N. Yonezawa, Y. Sato, H. Nishihara, K. Yamagata, H. Kimura; funding acquisition, Y. Sato, T. Kujirai, H. Kurumizaka, K. Yamagata, H. Kimura; supervision, H. Kurumizaka, K. Yamagata, H. Kimura.

---

**Figure S1.**
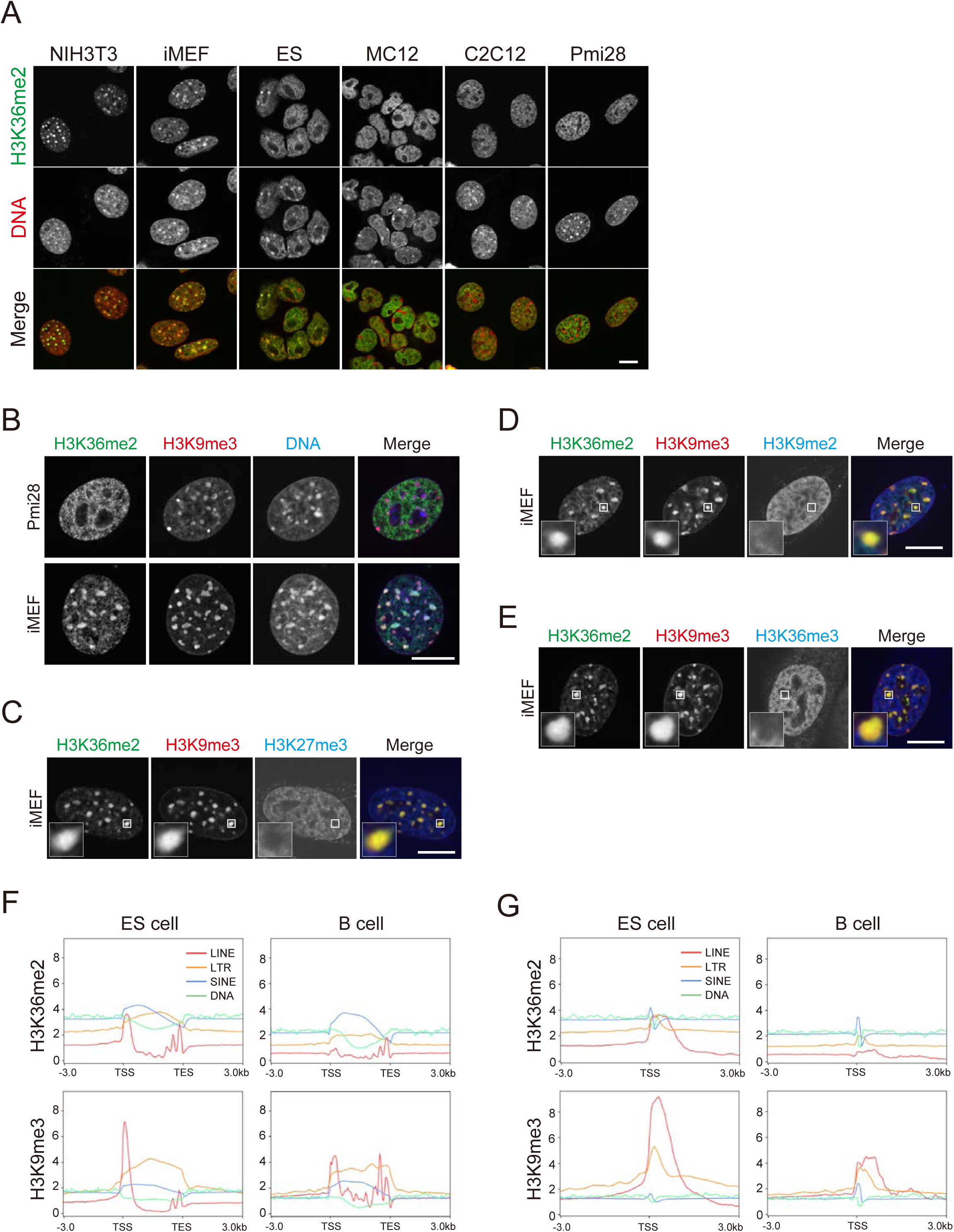
H3K36me2 is enriched in pericentromeric heterochromatin and LINE repeats in some mouse cell lines. (A-E) Immunofluorescence. Cells were fixed and stained with antibodies directed against histone modifications and Hoechst 33342. Single optical sections of confocal microscope images are shown. (A) H3K36me2 (green) and Hoechst33342 (red) in various mouse cell lines. (B) Pmi28 cells and iMEFs stained with another H3K36me2-specific antibody (green; rabbit monoclonal) with H3K9me3 (red) and Hoechst33342 (blue). (C-E) Localization of various modifications (blue), such as H3K27me3 (C), H3K9me2 (D), and H3K36me3 (E), respect to H3K36me2 (green) and H3K9me3 (red) in iMEFs. Insets show magnified views of the indicated PCH areas. (F and G) ChIP-seq signal enrichments of H3K36me2 and H3K9me3 on repeat elements in ESCs and B cells from the transcription start site (TSS) to the transcription end site (TES) (F) and around TSS (G). Scale bars, 10 μm.

**Figure S2.**
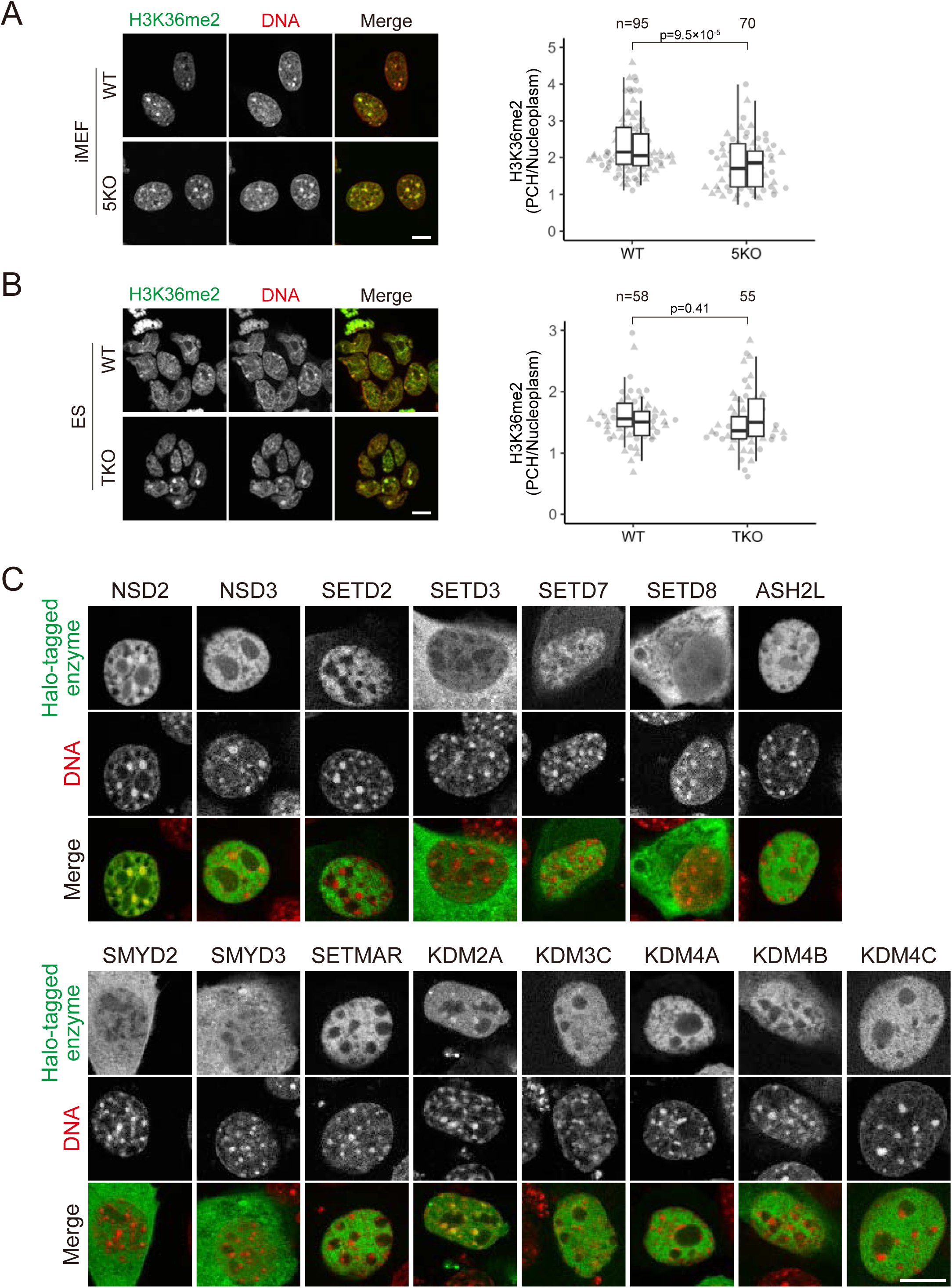
H3K36me2 PCH localization does not depend on H3K9me3 or DNA methylation and is induced by NSD2. (A and B) PCH localization of H3K36me2 in cells lacking H3K9me2/3 (A) or DNA methylation (B). 5KO iMEFs (lacking G9a, Glp, Setdb1, Suv39h1, and Suv39h2) (A) and TKO ESCs (lacking Dnmt1, Dnmt3a, and Dnmt3b) (B) were fixed and stained with anti-H3K36me2 (green) and Hoechst33342 (red). Single optical sections are shown on the left. Box plots display the intensity ratios of H3K36me2 in PCH to the nucleus. See Fig. 1 legend for the details of box plots. (C) Representative live-cell images of HaloTag-tagged H3K36 methylases and demethylases. NIH3T3 cells transfected with expression vectors for indicated Halo-tagged enzymes were stained with JF646 HaloTag ligand (green) and Hoechst33342 (red). Scale bars, 10 μm.

**Figure S3.**
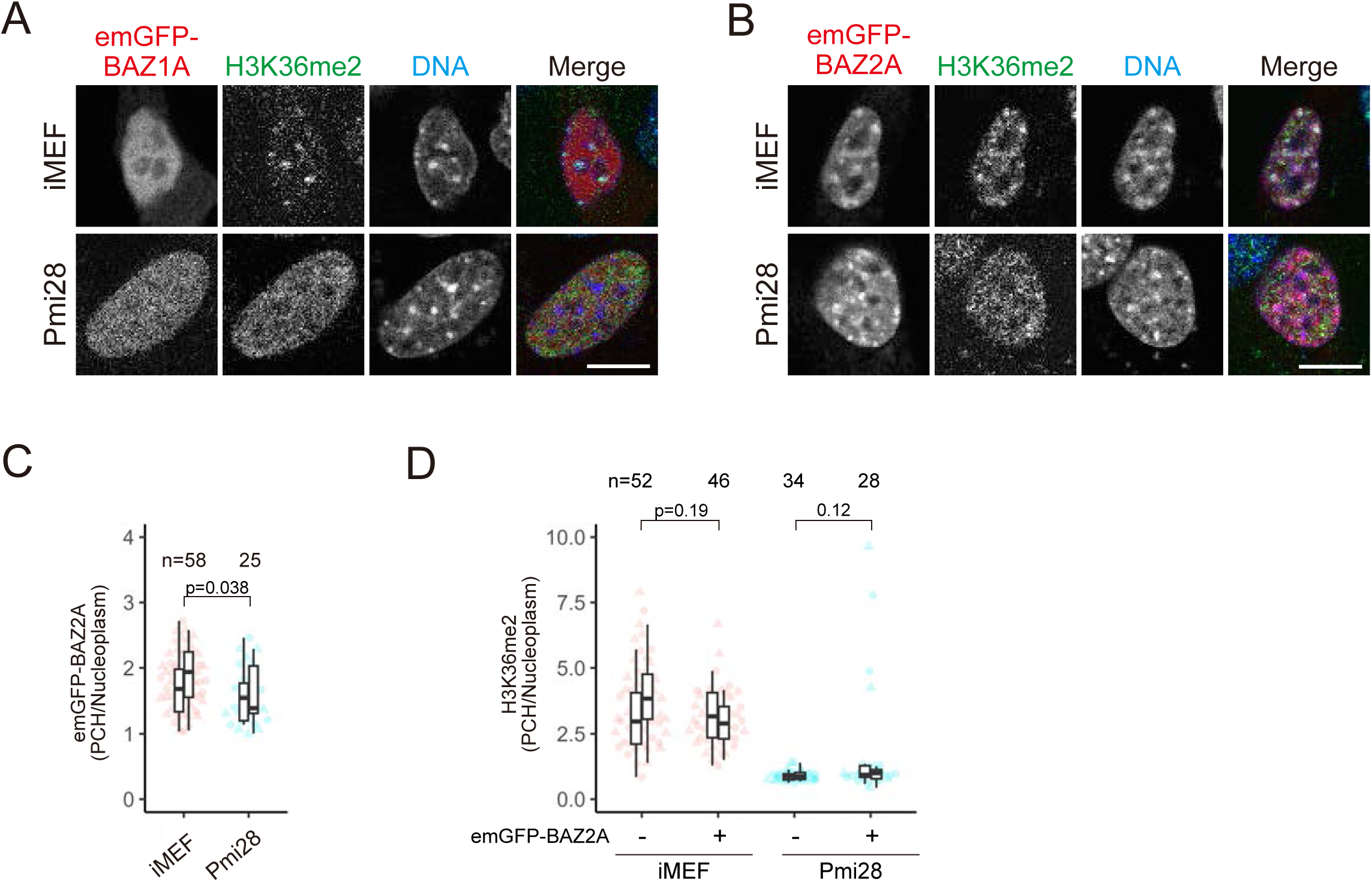
Effect of BAZ1A and BAZ2A on H3K36me2 localization. iMEFs and Pmi28 cells transfected with emGFP-BAZ1A (A) or emGFP-BAZ2A (B-D) expression vectors were stained with Hoechst 33342 for live-cell imaging (C). Cells were fixed and stained with anti-H3K36me2. (A and B) Examples of single optical sections of cells expressing emGFP-BAZ1A (A) and emGFP-BAZ2A (B) (red), H3K36me2 (green), and Hoechst33342 (blue). (C and D) PCH-to-nucleus intensity ratios of emGFP-BAZ2A in live cells (C) and H3K36me2 by immunofluorescence (D). See Fig. 1 legend for the details of box plots. Scale bars, 10 μm.

**Figure S4.**
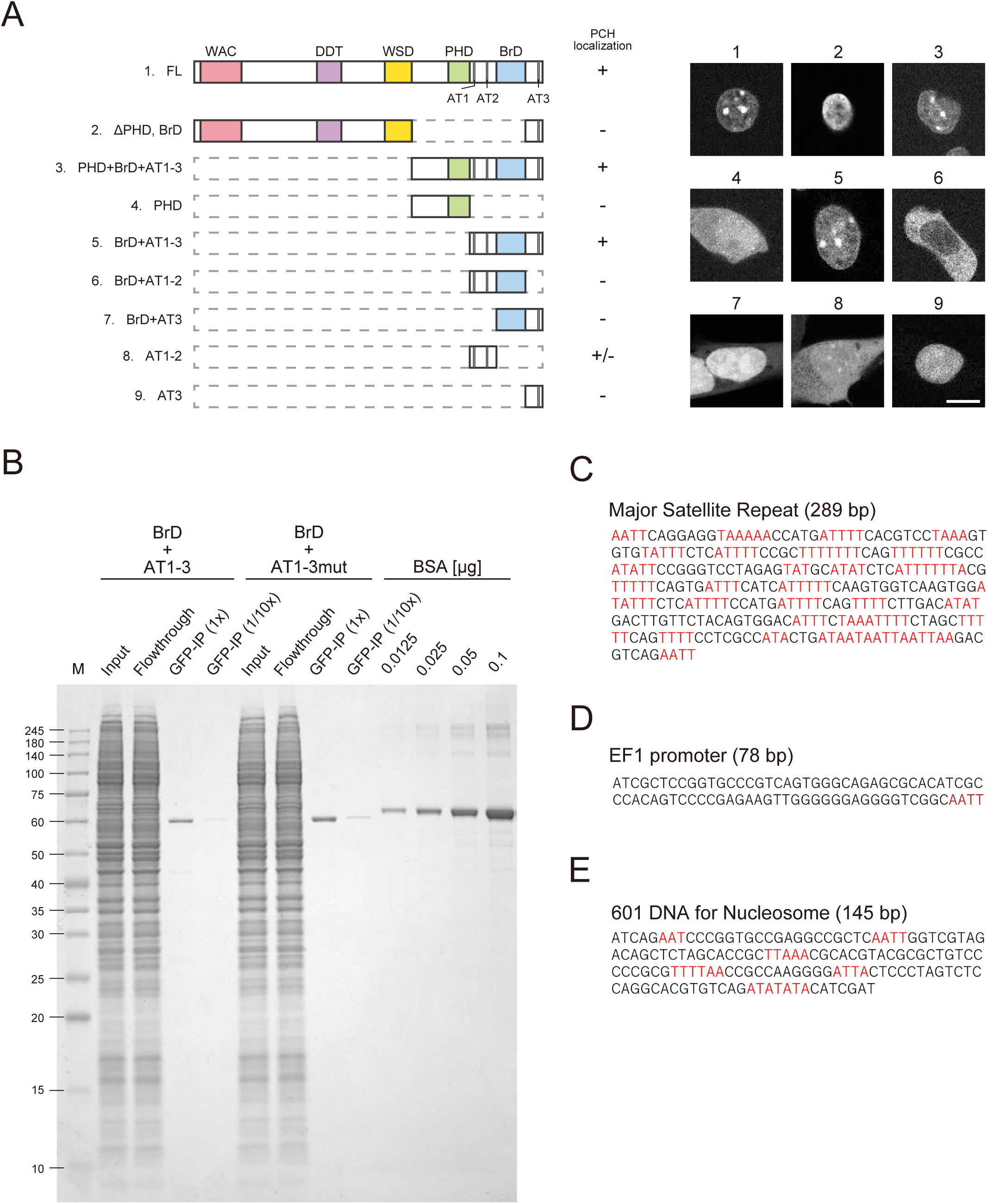
BAZ1B domain targeting PCH. (A) Schematic drawing of various truncated BAZ1B and their localizations. (B) Expression and purification of EGFP-tagged BAZ1B BrD+AT1-3 and BrD+AT1-3mut were evaluated by SDS-polyacrylamide gel electrophoresis and Coomassie Brilliant Blue staining. HEK293T cells transfected with the expression vectors were lysed and the supernatant (input) was incubated with GFP Trap anti-GFP nanobody magnetic beads at 4°C overnight. After removing the unbound fraction (Flowthrough), beads were washed, and aliquots were mixed with SDS-gel loading buffer (GFP-IP). Serial dilution series of bovine serum albumin (BSA) were loaded to estimate the amounts of recovered protein bound to beads. (C-E) Nucleotide sequences of DNA used for binding assay. (C) Major Satellite Repeats. (D) EF1 promoter. (E) 601 DNA. Sequences with more than three consecutive AT-bases are highlighted in red. Scale bar, 10 μm.

**Figure S5.**
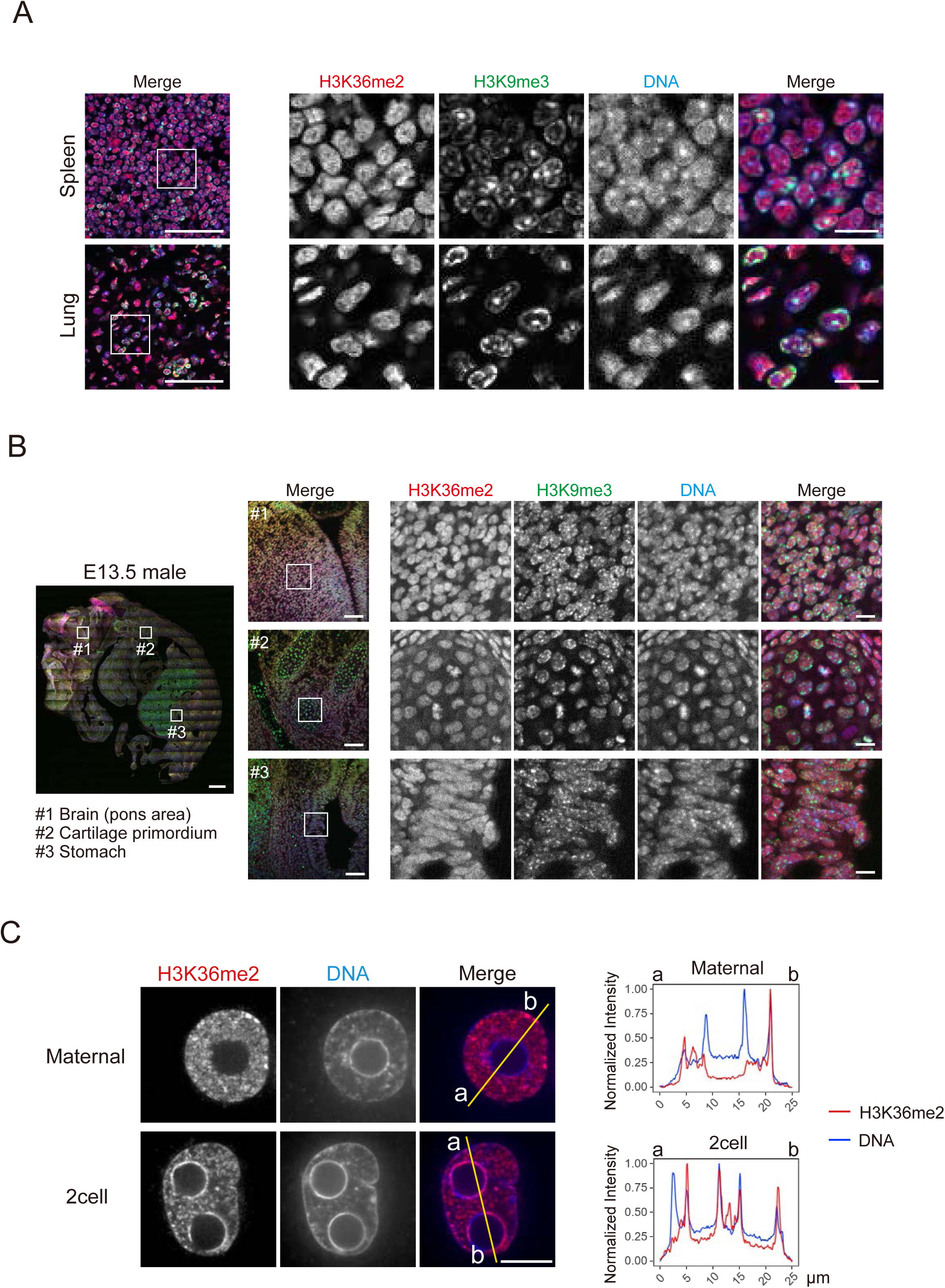
H3K36me2 in mouse tissues, E13.5, and preimplantation embryos. (A and B) Immunofluorescence of tissue sections (A) and 13.5-day embryo sections (B) stained with antibodies against H3K36me2 (red), H3K9me3 (green), and Hoechst 33342 (blue). Single optical sections of confocal microscope images are shown. (A) Low-power views of merged images are shown on the left (Scale bars, 50 μm). Individual and merged images of the indicated area are shown on the right (Scale bars, 10 μm). (B) A tiled image covering the whole embryo is shown on the left (Scale bar, 500 μm). Single images for the brain (pons area; #1), cartilage primordium (#2), and stomach (#3) (Scale bars, 50 μm) and magnified views of the indicated areas are shown from the middle to the right (Scale bars, 10 μm). (C) Different localization of H3K36me2 in maternal PN in zygotes and the 2-cell nucleus in preimplantation embryos analyzed by line plots. Intensity profiles of H3K36me2 and DNA along the yellow lines are shown. Scale bar, 10 μm.

**Table S1.**
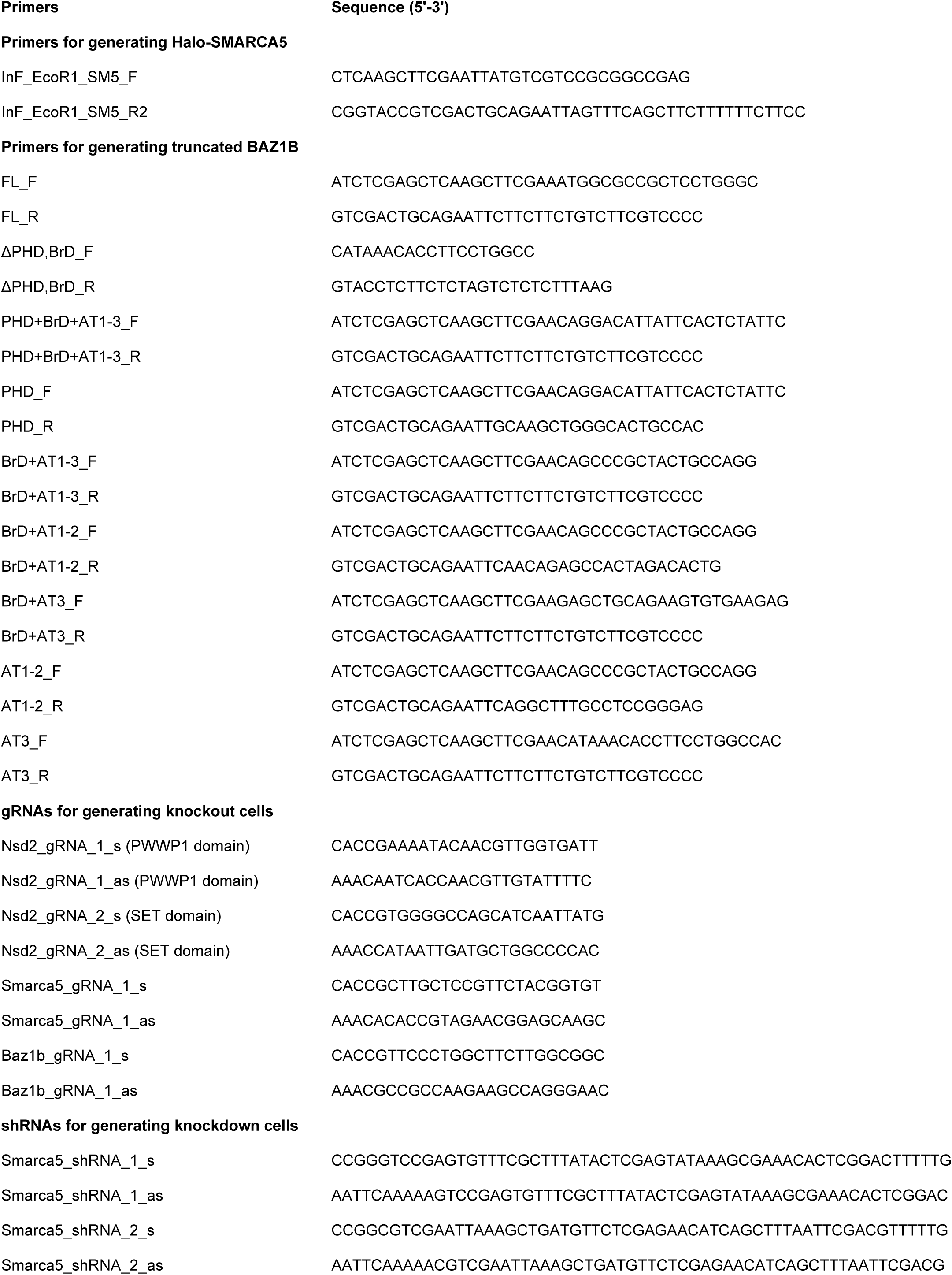
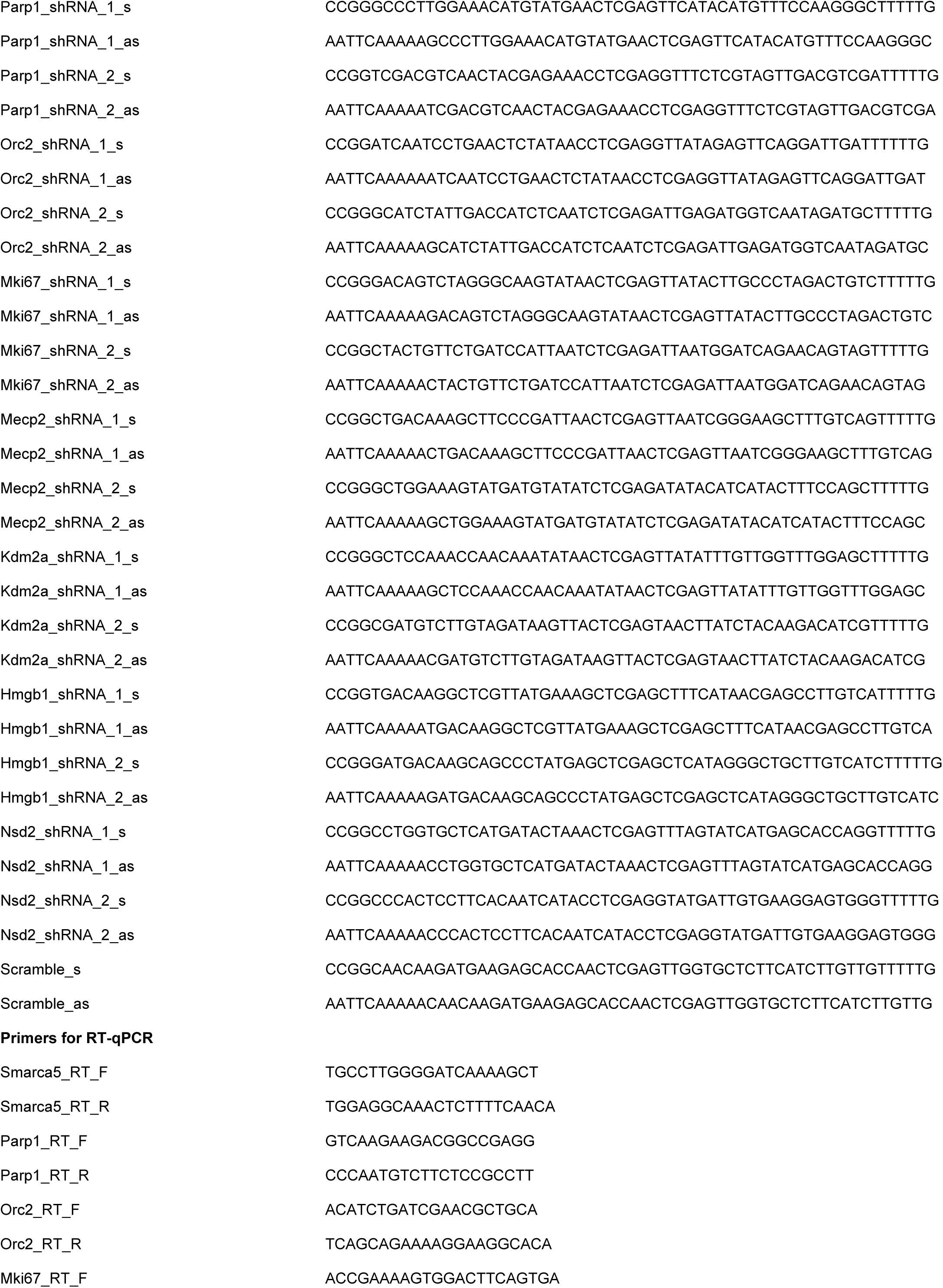
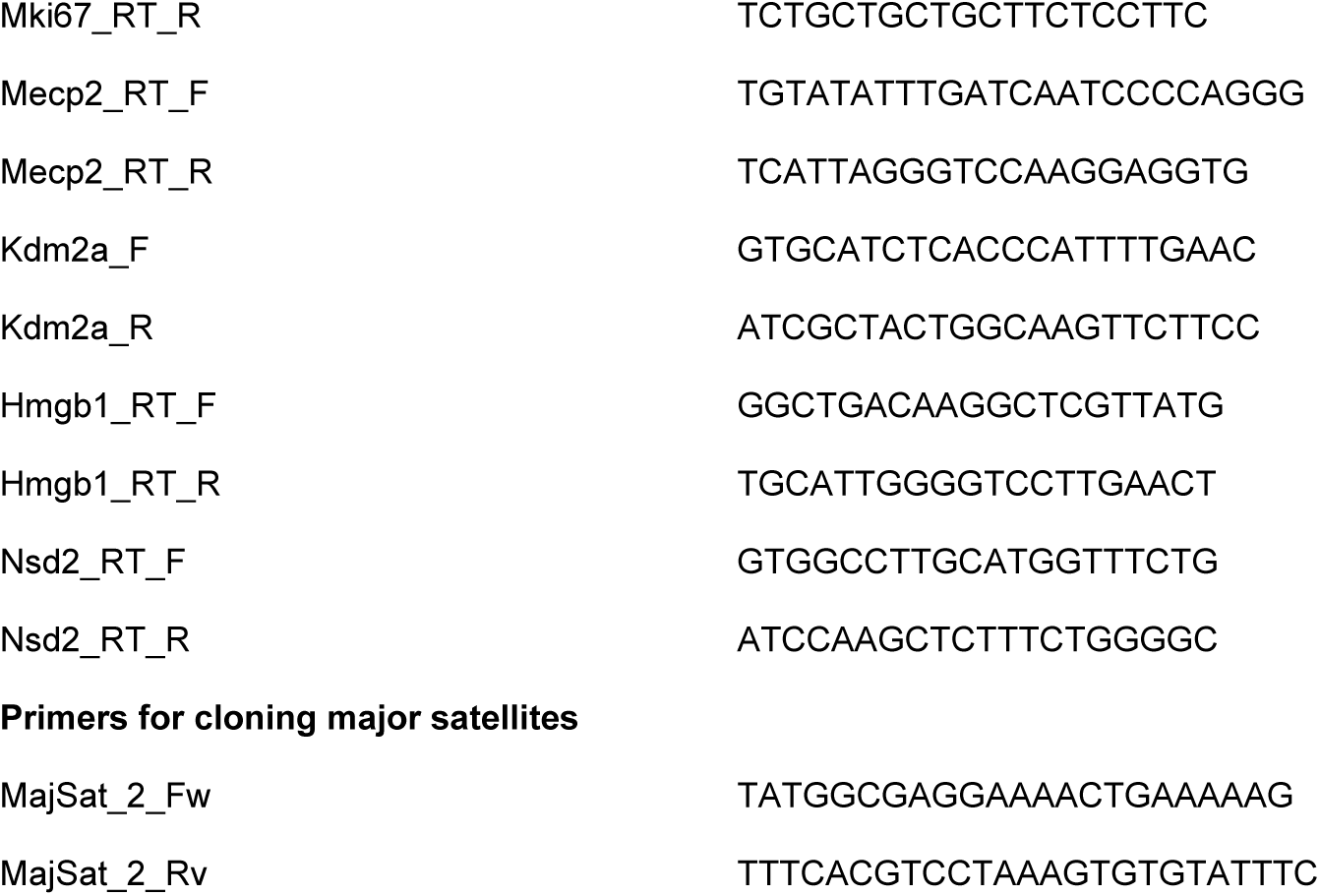
Oligonucleotides.

**Table S2.**
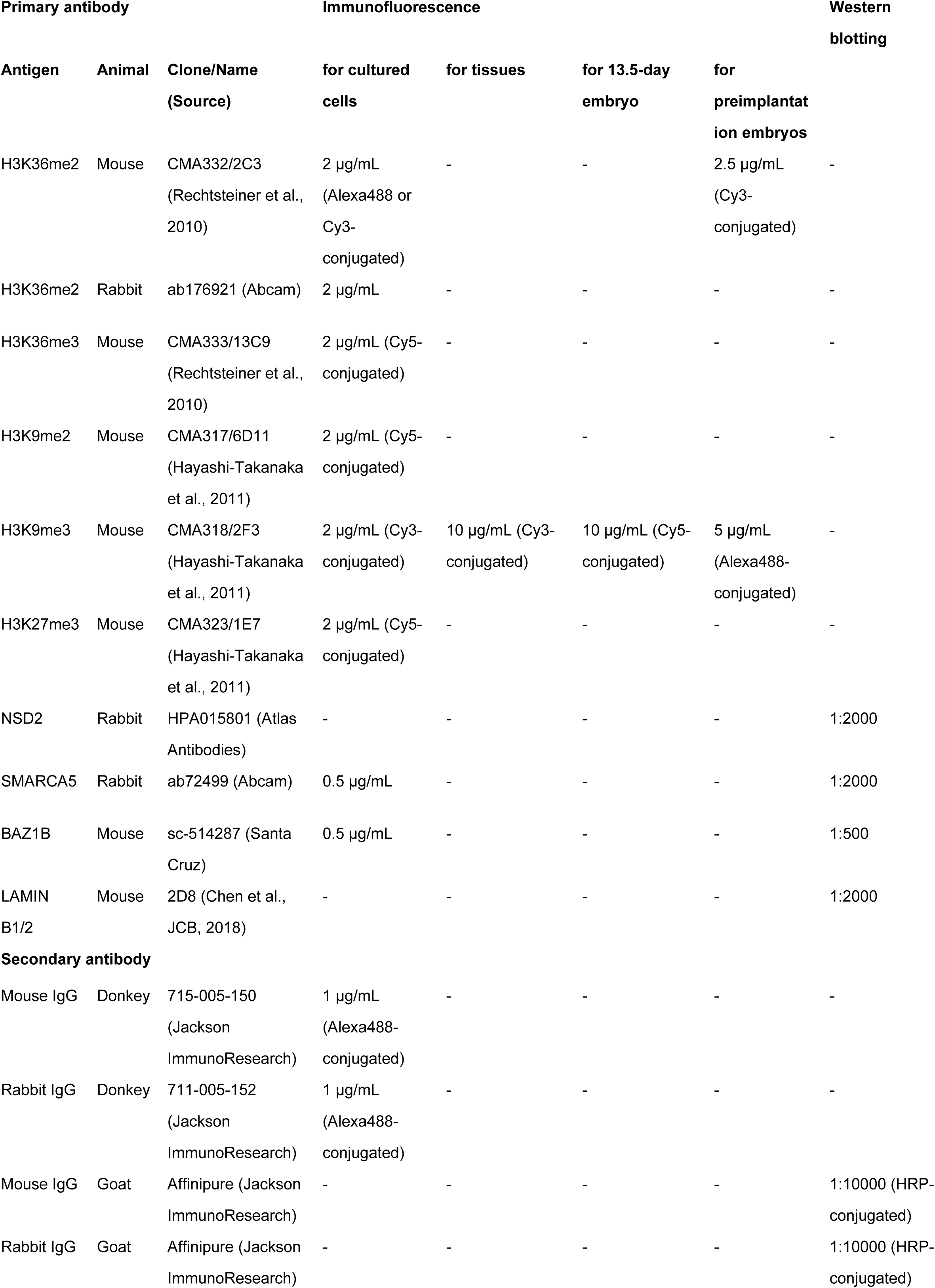
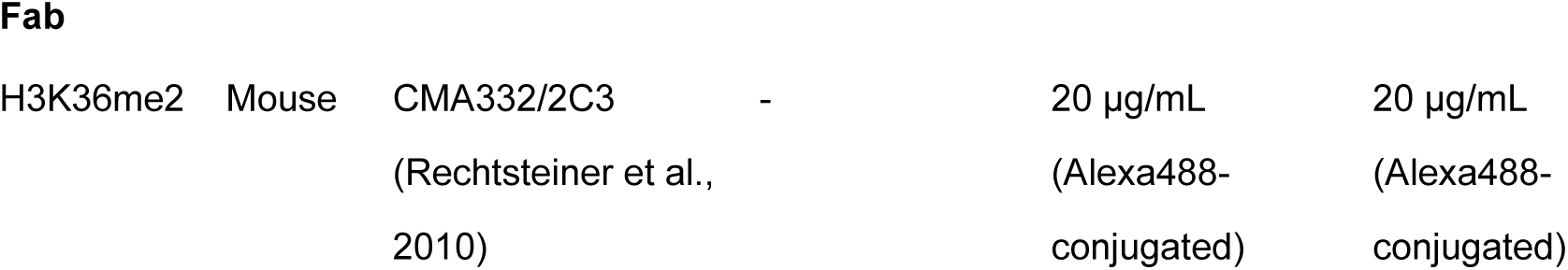
Antibodies.

## Notes

### Competing Interest Statement

The authors have declared no competing interest.

### Summary of Updates

A few minor errors were corrected and two references were added.

